# Mechanism and regulation of cargo entry into the Commander recycling pathway

**DOI:** 10.1101/2024.01.10.574988

**Authors:** Rebeka Butkovič, Alexander P. Walker, Michael D. Healy, Kerrie E. McNally, Meihan Liu, Kohji Kato, Brett M. Collins, Peter J. Cullen

## Abstract

Commander is a multiprotein complex that orchestrates endosomal recycling of diverse integral cargo proteins and in humans is required for normal skeletal, brain, kidney, and cardiovascular development. While the structure of this complex has recently been described, the central question of how cargo proteins are selected for entry into the Commander recycling pathway remains unclear. Here using recombinant protein reconstitution and *in silico* predictions we identify the evolutionary conserved mechanism through which the unstructured carboxy-terminal tail of the integral protein adaptor sorting nexin-17 (SNX17) directly binds to the Retriever sub-complex of Commander. SNX17 adopts an autoinhibited conformation where its carboxy-terminal tail occupies the cargo binding groove. Competitive cargo binding overcomes this autoinhibition, promoting SNX17 endosomal residency and the release of the carboxy tail for Retriever association. Using molecular cell biology and high-resolution microscopy, we establish the central importance of SNX17-Retriever association in the handover of integrin and lipoprotein receptor cargoes into pre-existing endosomal retrieval sub-domains for entry into the recycling pathway. In describing the principal mechanism of cargo entry into the Commander recycling pathway we provide key insight into the function and regulation of this evolutionary conserved sorting complex.

## INTRODUCTION

The intracellular eukaryotic endosomal network sorts and transports thousands of integral membrane proteins and their associated proteins and lipids (Cullen and Steinberg, 2018; Norris and Grant, 2020; Weeratunga et al., 2020). Central to network function are multi-protein complexes that coordinate sequence-dependent recognition of integral proteins with the biogenesis of vesicular and tubular transport carriers (McNally and Cullen, 2018; Chen et al., 2019; McDonald, 2021). Retriever is an essential cargo sorting complex and is a stable heterotrimer of VPS26C, VPS35L and VPS29, that together with the dodecameric CCDC22, CCDC93, COMMD (CCC) complex and DENND10, forms the 16-subunit Commander super-assembly (Phillips-Krawczak et al., 2015; Mallam and Marcotte, 2017; McNally et al., 2017; Boesch et al., 2023; Healy et al., 2023; Laulumaa et al., 2023). Defects in Commander assembly and function are associated with metabolic disorders including hypercholesterolemia (Bartuzi et al., 2016; Fedoseienko et al., 2018; Rimbert et al., 2020; Vos et al., 2023), viral infection (Daniloski et al., 2021; Zhu et al., 2021), and lead to Ritscher-Schinzel syndrome, a multi-system developmental disorder characterized by abnormal craniofacial features, cerebellar hypoplasia, and stunted cardiovascular development (Kolanczyk et al., 2015; Kato et al., 2020; Otsuji et al., 2023). In the Commander trafficking pathway sequence-dependent integral protein recognition is principally mediated by the cargo adaptor sorting nexin-17 (SNX17) (McNally et al., 2017), the FERM domain of which binds to a ØxNxx[Y/F] sorting motif presented in the cytoplasmic facing domains of integral proteins (where Ø is a hydrophobic residue and x is any residue) (van Kerkhof et al., 2005; Bottcher et al., 2012; Steinberg et al., 2012; Ghai et al., 2013). Over 100 integral proteins require SNX17 and Retriever for their endosomal sorting through the Commander axis including members of the integrin and lipoprotein receptor families (McNally et al., 2017). Fundamental to our understanding of the Commander pathway and the dissection of its functional role in health and disease is a central unanswered question: how is SNX17 coupled to Retriever to allow access into the Commander endosomal retrieval and recycling pathway and how is this coupling regulated?

## RESULTS AND DISCUSSION

### SNX17 associates with Commander via its extended C-terminal domain

Previously we showed that the carboxy-terminal unstructured ^465^IGDEDL^470^ tail of SNX17, Leu470 being the terminal residue, can bind directly to the PDZ domain of the PDLIM family of proteins (Healy et al., 2022). The same sequence is essential for binding to Commander, and we speculated that this was through direct binding to Retriever (McNally et al., 2017; Healy et al., 2022). To test this association, we used biGbac insect cell expression (Healy et al., 2023) to purify human Retriever and full-length human SNX17 (Supp. Fig. 1A). When SNX17 was incubated with nickel affinity resin bound Retriever we observed specific but non-stoichiometric association (Fig. 1A). Consistent with the requirement of the carboxy-terminal Leu470 residue (McNally et al., 2017), recombinant SNX17(L470G) bound to Retriever at significantly lower levels (Fig. 1A, Supp. Fig. 1B). These data establish that SNX17 directly binds to Retriever through a mechanism that involves its unstructured carboxy-terminal tail, a region that is highly conserved across eukaryotic SNX17 (Supp. Fig. 1C).

**Figure 1.**
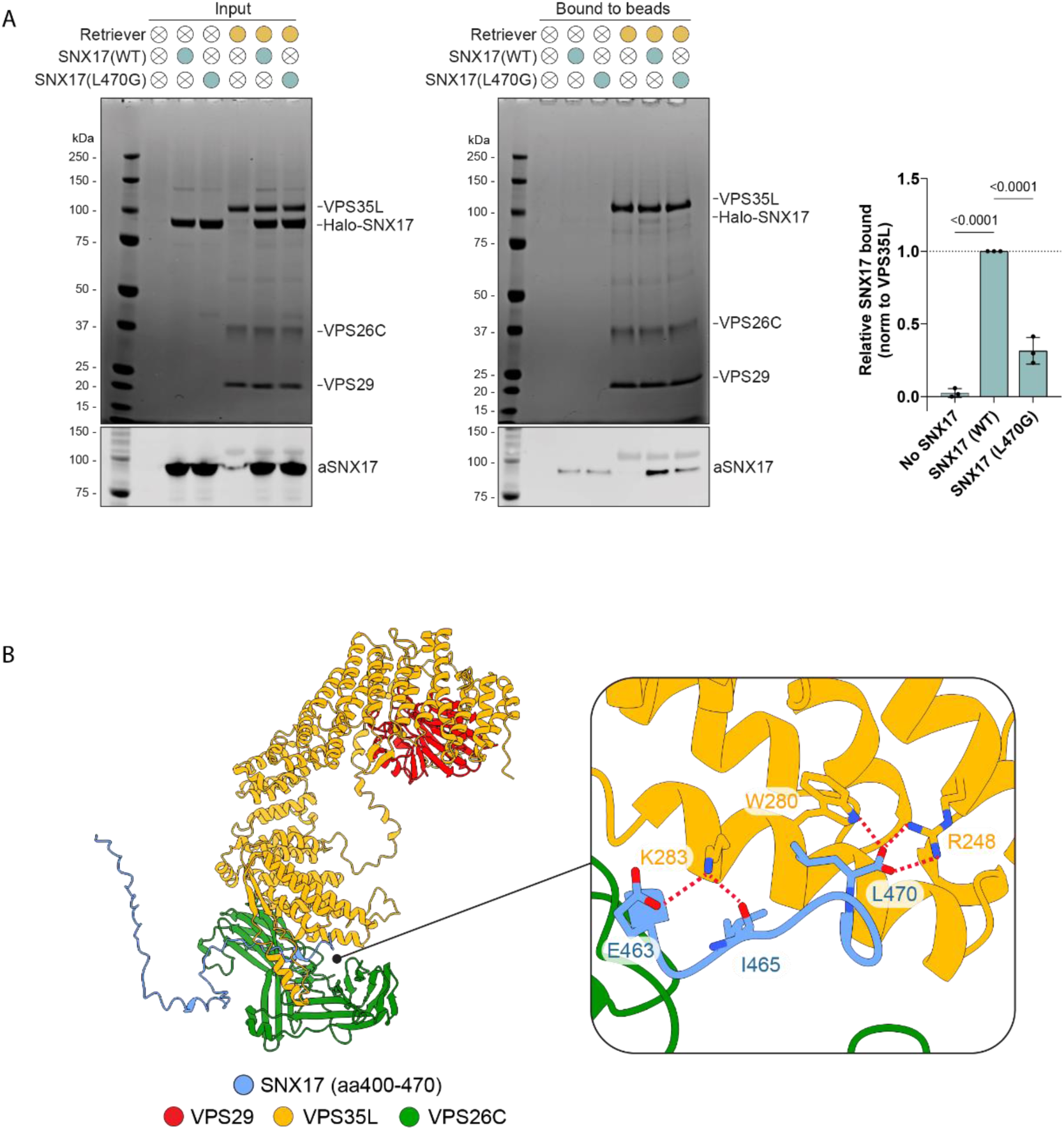
SNX17 directly binds to the Retriever complex. **(A)** Purified His-tagged Retriever was mixed with purified SNX17 (WT) or SNX17 (L470G) and incubated with anti-His-tag TALON® Superflow beads. Input mixtures (left) and protein bound to the beads after washing (middle) were analysed by SDS-PAGE followed by Coomassie staining and western blotting. SNX17 bound to the beads was quantified and normalised to the level of VPS35L (right). n = 3, 1-way ANOVA with Dunnett’s multiple comparison test, error bars represent s.d. **(B)** AlphaFold2 predictions show a high confidence interaction between the unstructured carboxy-terminal region of SNX17 and Retriever (Fig. S4A).

Retriever is assembled around a central VPS35L subunit to which VPS26C and VPS29 associate at spatially distant amino- and carboxy-terminal regions of the VPS35L a-solenoid (Fig. 1B) (Boesch et al., 2023; Healy et al., 2023; Laulumaa et al., 2023). We employed AlphaFold2 modelling (Jumper et al., 2021; Mirdita et al., 2022) to predict the association between Retriever and the unstructured human SNX17 tail corresponding to residues Gly400-to-Leu470. This predicted a high confidence model where the ^465^IGDEDL^470^ motif of SNX17 bound to a pocket in VPS35L defined by Arg248, Trp280, and Lys283, that resided close to the VPS35L:VPS26C interface: this pocket is highly conserved across eukaryotic Retriever (Fig. 1B). The interface between SNX17 and Retriever extended to include upstream residues ^459^NFAF^462^ in the SNX17 tail engaging primarily with VPS26C. One striking feature of the predicted complex is that the extreme carboxy-terminal Leu470 residue of SNX17 makes extensive contact with VPS35L through hydrophobic interaction with Trp280 and via an electrostatic interaction of the carboxy-terminal carboxyl group with Arg248 (Fig. 1B). This agrees with the critical importance of Leu470 for interaction with Retriever (Fig. 1) and the larger Commander complex in cells (McNally et al., 2017; Healy et al., 2022).

Consistent with SNX17 binding to Retriever being a feature of the VPS35L:VPS26C interface, a VPS35L mutant that specifically disrupted binding to VPS26C, VPS35L(R293E) (Healy et al., 2023), failed to associate with SNX17 in quantitative cell-based immunoprecipitations where mCherry-SNX17 was co-expressed alongside VPS35L-GFP (Fig. 2A). In contrast a mutant that selectively disrupted VPS29 binding to VPS35L, VPS35L(L35D) (Healy et al., 2023), had no effect on SNX17 association (Fig. 2A). Based on the predicted structure of the SNX17-Retriever complex we performed mutagenesis of the proposed SNX17 binding pocket. These mutations confirmed the AlphaFold2 model; VPS35L(R248A), -(W280A), and -(K283E) all showed a pronounced loss of SNX17 association (Fig. 2B). Highlighting the selectivity of these mutations, all VPS35L mutants targeting SNX17 binding retained assembly into Retriever, binding to the CCC complex, and assembly into Commander.

**Figure 2.**
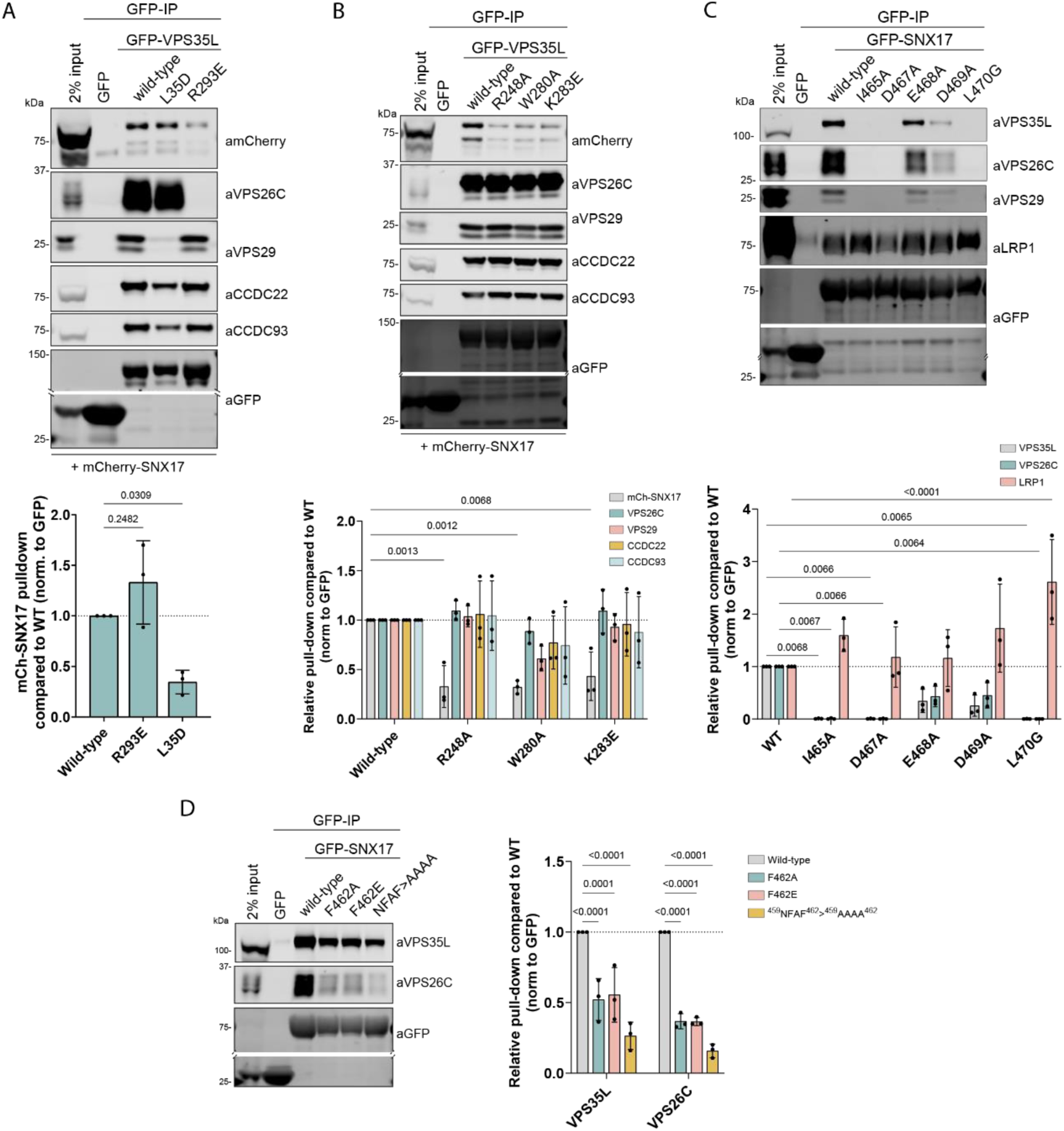
SNX17 binds to the VPS35L-VPS26C interface of Retriever complex. **(A)** HEK293T cells were transiently co-transfected with mCherry-SNX17 and either GFP, VPS35L-GFP or VPS35L-GFP mutants that perturb VPS35L-VPS26C (VPS35L(R293E)) and VPS35L-VPS29 (VPS35L(L35D)) associations prior to GFP-nanotrap isolation and quantitative western analysis of protein band intensities. n = 3, 1-way ANOVA with Dunnett’s multiple comparison test, error bars represent s.d. **(B)** HEK293T cells were transiently co-transfected with GFP, or VPS35L-GFP or VPS35L-GFP mutants that target SNX17 binding, and mCherry-SNX17. Protein lysates were then used in GFP-nanotrap experiments. Below, the quantitative analysis of protein band intensities is shown. n = 3, 2-way ANOVA with Dunnett’s multiple comparison test, error bars represent s.d., only changes with p < 0.05 are shown. **(C, D)** HEK293T cells were transiently co-transfected with GFP or GFP-SNX17 or GFP-SNX17 mutants in all conserved residues of the terminal ^465^IGDEDL^470^ motif (C) or conserved ^459^NFAF^462^ motif (D) that target Retriever binding. Protein lysates were then used in GFP-nanotrap experiments. Below, the quantitative analysis of protein band intensities is shown. n = 3, 2-way ANOVA with Dunnett’s multiple comparison test, error bars represent s.d., only changes with p < 0.05 are shown.

To further validate the AlphaFold2 model, we performed targeted mutagenesis of the SNX17 ^459^NFAF^462^ and ^465^IGDEDL^470^ sequences. Quantitative GFP-trap experiments in transiently transfected HEK293T cells established that GFP-SNX17(I465A), -(D467A) and -(L470G) mutants led to near-complete loss of Retriever binding (Fig. 2C). Similarly, SNX17(NFAF-AAAA) and the more conservative SNX17(F462A) and -(F462E) mutants also displayed reduced Retriever and CCC complex association (Fig. 2D). In contrast, SNX17(E468A) and -(D469A) mutations had only a modest impact on Retriever association (Fig. 2C), consistent with the predicted structure where these sidechains make no direct contacts with the Retriever complex (Fig. 1B). Collectively these data show that Retriever coupling of SNX17 is mediated by motifs within its unstructured carboxy termini binding to surfaces at the VPS26C:VPS35L interface of Retriever. Modeling the association of Retriever and Snx17 from *Drosophila melanogaster* (Supp. Fig. 2B) and other species such as zebrafish (not shown) predict essentially identical interactions between SNX17 and the Retriever assembly, supporting the evolutionary conservation of the coupling mechanism.

Although the molecular details are all very different, at a general level this coupling mechanism is reminiscent of how the related Retromer complex engages its own cargo adaptors sorting nexin-3 (SNX3) and sorting nexin-27 (SNX27): an unstructured region of SNX3 binding to the VPS26A/B:VPS35 interface (Lucas et al., 2016; McGough et al., 2018; Leneva et al., 2021) and the PDZ domain of SNX27 binding to VPS26A/B (Steinberg et al., 2013; Gallon et al., 2014) (Supp. Fig. 2A). While the relative orientation of Retriever to the endosomal membrane has yet to be resolved, we speculate that the similarity in SNX3-Retromer binding, and the ability of SNX17 to associate with PI(3)P, indicates that the SNX17 binding VPS26C-VPS35L interface likely lies in close proximity to the membrane surface (Supp. Fig. 2A).

### The SNX17-Commander interaction is required for cargo recycling to the plasma membrane

To test the functional importance of direct SNX17-Retriever coupling we performed rescue experiments in a VPS35L CRISPR/Cas9 knock-out RPE1 cell line. Re-expressed wild-type GFP-VPS35L localised to the endosomal network as expected, as did the VPS35L(R248A) mutant which associates normally to Retriever and the CCC complex but is defective in binding to SNX17 (Fig. 3A). Previous studies have shown that loss of VPS35L expression leads to Commander dysfunction including a reduction in the association of endogenous COMMD1, a marker of the CCC complex and Commander assembly to Retromer labelled endosomes (Healy et al., 2023). This phenotype was rescued by re-expression of either GFP-VPS35L or GFP-VPS35L(R248A), although there was a slight but significant trend towards reduced COMMD1 recruitment for the R248A mutant (Fig. 3A). At the functional level, VPS35L KO leads to a reduction in the steady-state cell surface enrichment of a5b1-integrin and the mis-sorting of the internalized integrin into LAMP1-positive late endosomes/lysosomes (McNally et al., 2017), but does not significantly perturb whole-cell levels of SNX17 (Supp. Fig. 2C). In imaging experiments these phenotypes were fully rescued by re-expression of wild-type GFP-VPS35L but not by the SNX17-binding defective GFP-VPS35L(R248A) mutant (Fig. 3B). Quantification of the level of cell surface a5b1-integrin confirmed the plasma membrane recycling defect caused by SNX17 binding deficiency and extended the significance of the coupling mechanism to another functionally important SNX17 cargo the lipoprotein receptor LRP1 (van Kerkhof et al., 2005) (Fig. 3C). Together, these data reveal the core mechanism of SNX17 coupling to Retriever and the essential importance of coupling for endosomal retrieval and recycling of SNX17 selected cargoes through the Commander pathway.

**Figure 3.**
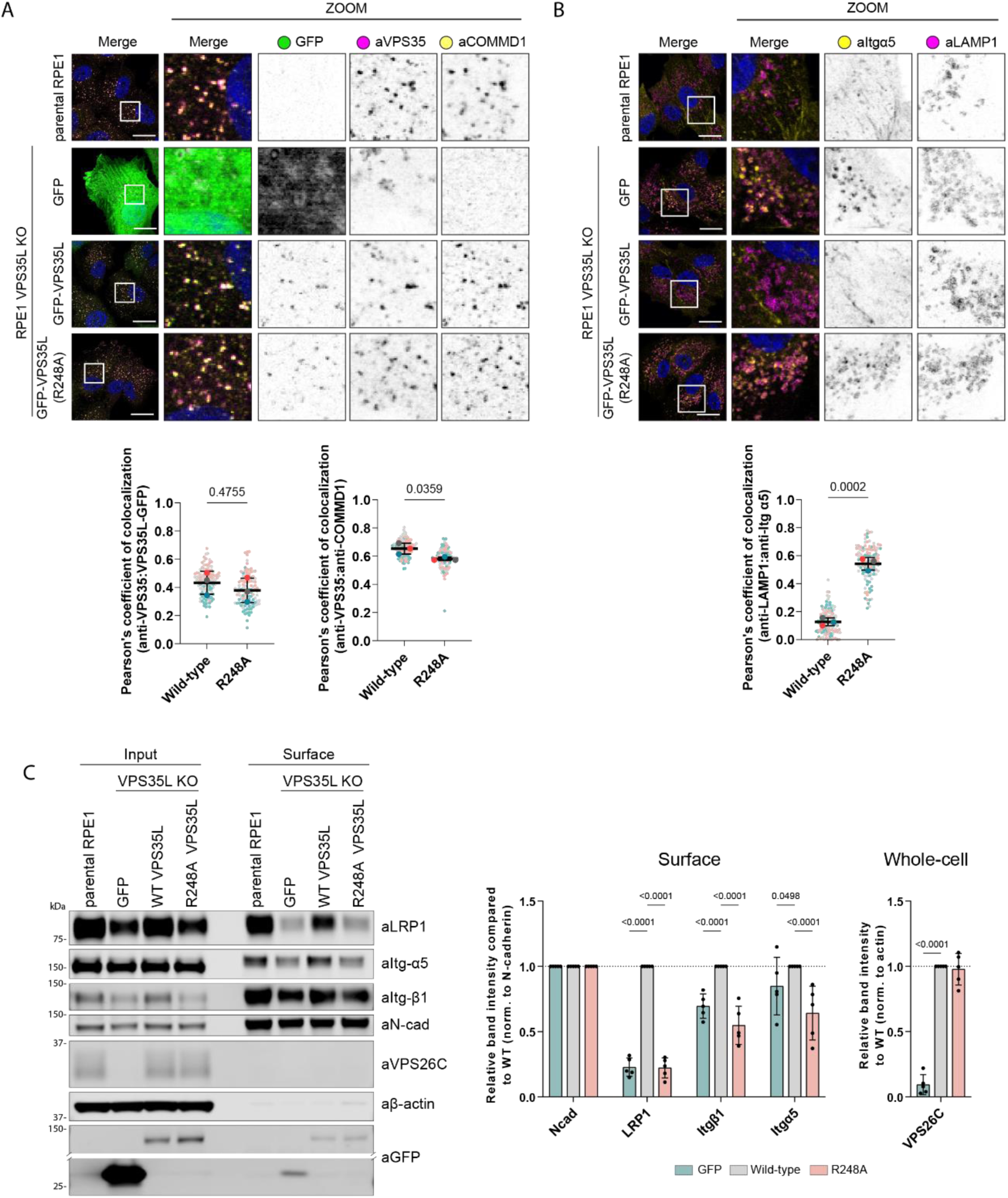
SNX17-Retriver coupling is essential for Retriever-cargo retrieval to plasma membrane. **(A)** VPS35L KO RPE1 cells were lentivirally transduced with GFP, VPS35L-GFP or VPS35L-GFP(R248A). The stably expressed VPS35L(R248A) localizes to endosomes and can partially rescue COMMD1 localization. Scale bars corresponds to 20 µm. Pearson’s coefficients were quantified from 3 independent experiments (GFP-VPS35 coloc. wt: n=114 cells, R248A: n=107 cells, VPS35-COMMD1 coloc. wt: n=117 cells, R248A: n=106 cells). Pearson’s coefficients for individual cells and means are presented by smaller and larger circles, respectively, colored according to the independent experiment. The means (n = 3) were compared using a two-tailed unpaired t-test. Error bars represent the mean, S.D. **(B)** VPS35L KO RPE1 cells were lentivirally transduced with GFP, VPS35L-GFP or VPS35L-GFP(R248A). The stably expressed VPS35L(R248A) failed to rescue Itgα5 missorting as evidenced by increased co-localisation with lysosomal marker LAMP1. Scale bars corresponds to 20 µm. Pearson’s coefficients were quantified from 3 independent experiments (wt: n=125 cells, R248A: n=113 cells). Pearson’s coefficients for individual cells and means are presented by smaller and larger circles, respectively, colored according to the independent experiment. The means (n = 3) were compared using a two-tailed unpaired t-test. Error bars represent the mean, s.d. **(C)** Cell surface proteins were biotinylated in stably rescued RPE1 cells and enriched with streptavidin pull-down to analyze surface protein levels. The VPS35L(R248A) mutant failed to rescue cell surface levels of Itgα5, Itgβ1 and LRP1, but stabilised VPS26C. The quantitative analysis of protein band intensities is shown. The band intensities were normalised to the respective cell surface N-cadherin levels. n = 5, 2-way ANOVA with Dunnett’s multiple comparison test, error bars represent s.d., only changes with p < 0.05 are shown.

### A SNX17 autoinhibitory sequence regulates cargo and Retriever interactions

We next explored how the coupling between SNX17 and Retriever could be regulated. The FERM domain of SNX17 comprises three submodules, F1, F2 and F3 (Ghai et al., 2013). Alphafold2 modelling suggested that cargo proteins carrying the conserved ØxNxx[Y/F] sorting motif, including β1-integrin and LRP1, bind through β-sheet augmentation in a complementary groove of the F3 module, in agreement with previous X-ray crystallographic structures (e.g. P-selectin (PDB:4GXB) (Fig. 4A, D). In models of apo SNX17 from various species, the carboxy-terminal ^459^NFAF^462^ sequence (human numbering) invariably formed an intramolecular association with the F3 groove, mimicking cargo sequences in what would be an autoinhibitory conformation (Fig. 4A). The carboxy-terminal ^465^IGDEDL^470^ also adopted an intramolecular conformation that would be mutually exclusive of Retriever binding. In our initial modelling of the SNX17 interaction with Retriever we used only the isolated carboxy-terminal disordered region, which yields essentially identical structure predictions across all five models (Fig. 1B, Supp. Fig 4A). However, when we expanded the AlphaFold2 modelling to incorporate full-length SNX17 only two of the five predicted structures displayed the same mechanism of SNX17 binding to the VPS26C:VPS35L interface. Three of the five predicted structures showed no interaction with Retriever; in these cases, the ^459^NFAF^462^ sequence of SNX17 adopted the intramolecular association as seen in apo SNX17 predictions (Fig. 4A; Supp. Fig. 2D). This led us to speculate that such a conformation may reflect an autoinhibited state that negatively regulates coupling to Retriever and hypothesized that such autoinhibition could be released by competitive binding of ØxNxx[Y/F]-containing cargo.

**Figure 4.**
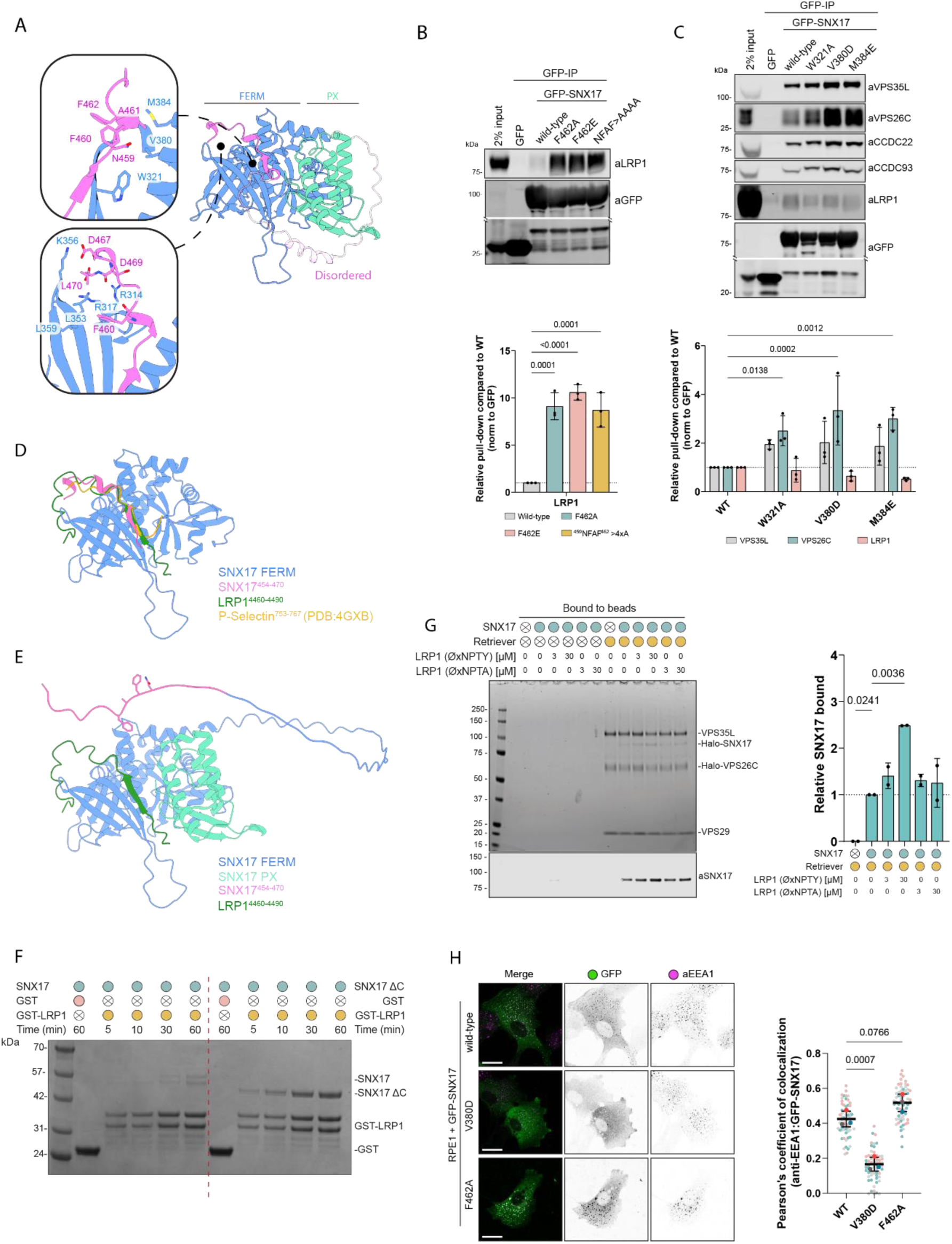
Intramolecular association between SNX17 FERM domain and ^459^NFAF^462^ autoinhibits SNX17 binding to cargo and Retriever. **(A)** The C-terminus of SNX17 contains an ^459^NFAF^462^ motif highlighted in bright pink that is predicted by AlphaFold2 (Fig. S4B) to bind into the canonical cargo binding pocket, in addition a number of residues in the extreme C-terminus (^467^DEDL^470^) are predicted to stabilize this interaction. **(B)** HEK293T cells were transiently co-transfected with GFP, or GFP-SNX17 or GFP-SNX17 mutants in the ^459^NFAF^462^ motif to target its intra-molecular association with SNX17-FERM domain. Protein lysates were then used in GFP-nanotrap experiments. Below, the quantitative analysis of protein band intensities is shown. n = 3, 1-way ANOVA with Dunnett’s multiple comparison test, error bars represent s.d. **(C)** HEK293T cells were transiently co-transfected with GFP, or GFP-SNX17 or GFP-SNX17 mutants in the FERM(F3) domain to target its intra-molecular association with the^459^NFAF^462^ motif. Protein lysates were then used in GFP-nanotrap experiments. Below, the quantitative analysis of protein band intensities is shown. n = 3, 1-way ANOVA with Dunnett’s multiple comparison test, error bars represent s.d., only changes with p < 0.05 are shown. **(D)** Overlay of the FERM domain of SNX17 bound to P-selectin (PDB: 4GXB), LRP1 (Fig. S4C) and the intramolecular SNX17 peptide (Fig. S4B) (as predicted by AlphaFold2) shows a clear overlap in peptide occupancy within the canonical cargo binding pocket. **(E)** When full length SNX17 is modeled with the cytoplasmic tail of LRP1^4460-4490^ the intramolecular interaction is perturbed, LRP1 preferentially binds the cargo binding pocket which in turn releases the disordered carboxy-tail of SNX17. **(F)** A GST tagged fragment of the LRP1 cytoplasmic tail was mixed with purified SNX17 (WT) (left) or SNX17^1-390^ (ΔC) (right) for various lengths of time before been washed and analyzed via Commassie staining of an SDS-PAGE gel. Removal of the disordered carboxy-tail increased both the rate and degree of SNX17 recruitment to GST-LRP1 consistent with the proposed autoinhibitory model. GST was used as a negative control to confirm the specificity of the interaction. **(G)** Purified His-tagged Retriever was mixed with purified SNX17 (WT) and 3 µM - 30 µM of LRP1 (ØxNPTY) or LRP1 (ØxNPTA) peptide. The mixtures were incubated with anti-His-tag TALON® Superflow beads, then input mixtures and protein bound to the beads after washing were analyzed by SDS-PAGE followed by Coomassie staining and western blotting. SNX17 bound to beads was quantified and normalised to the level of VPS35L (right). n = 2, 1-way ANOVA with Dunnett’s multiple comparison test, error bars represent s.d., only changes with p < 0.05 are shown. **(H)** RPE1 cells were transiently transfected with GFP-SNX17 wild-type, or SNX17(V380D) or SNX17(F462A) mutants that decrease or enhance cargo binding, respectively. Fixed cells were examined with confocal microscope, and the localisation of the GFP-tagged constructs was compared to the localisation of early endosome marker EEA1. Scale bars corresponds to 20 µm. Pearson’s coefficients were quantified from 3 independent experiments (wt: n=62 cells, V380D: n=60 cells, F462A n=64 cells). Pearson’s coefficients for individual cells and means are presented by smaller and larger circles, respectively, coloured according to the independent experiment. The means (n = 3) were compared using a were compared using a 1-way ANOVA with Dunnett’s multiple comparison test. Error bars represent the mean, s.d.

To test this hypothesis, we quantified the association of LRP1, a model ØxNxx[Y/F] cargo, to SNX17 wild-type and mutants targeting the conserved (Supp. Fig. 1C) intramolecular ^459^NFAF^462^ motif: SNX17(NFAF-AAAA), SNX17(F462A) and -(F462E). In all cases, immuno-isolation of GFP-tagged SNX17 mutants from HEK293T cells revealed a robust enhancement of LRP1 binding compared to the wild-type protein (Fig. 4B). This suggests that weakening or ablating the intramolecular interaction of the ^459^NFAF^462^ sequence allows for increased intermolecular binding of ØxNxx[Y/F]-containing cargo to the FERM domain. We modelled the association of full-length SNX17 with the cytosolic tail of LRP1 using AlphaFold2 which showed the expected β-sheet augmentation of the LRP1 ^4470^NPTY^4473^ sequence with the FERM domain (Ghai et al., 2013) and a displacement of the intramolecular ^459^NFAF^462^ interaction (Fig. 4E).

The mutations in the SNX17 carboxy-terminus that enhance LRP1 binding also perturb intermolecular binding of the SNX17 carboxy-terminus to Retriever (Fig. 2D, 4B), therefore we designed a set of complementary mutations in the SNX17 FERM domain required for binding to ØxNxx[Y/F]-containing cargo (Ghai et al., 2013); we predicted that these would relieve the autoinhibition with the carboxy-terminal ^459^NFAF^462^ sequence and thus enhance Retriever association. Indeed, the GFP-tagged SNX17(W321A), -(V380D) and -(M384E) mutants all displayed a modest but significant increase in binding to Retriever and the Commander super-assembly (Fig. 4C). These mutants also showed a reduced binding to the LRP1 cargo, consistent with binding being mediated through the same groove.

To demonstrate the presence of an autoinhibitory interaction we next preformed a GST pull down with recombinant wild-type SNX17 and SNX17ΔC, a deletion mutant lacking the carboxy-terminal tail motifs predicted to form the intramolecular inhibition. Consistent with the autoinhibitory model, recruitment of SNX17ΔC to a recombinant GST-fusion of the cytoplasmic tail of LRP1 was significantly higher than observed to wild-type SNX17 (Fig. 4F, Supp. Fig. 2E). Interestingly, at 60 min of co-incubation low level recruitment of wild-type SNX17 to GST-LRP1 was observed consistent with this peptide being sufficient to displace the SNX17 intramolecular autoinhibition.

Finally, to directly test that cargo-binding can relieve auto-inhibition and promote subsequent Retriever binding, we purified recombinant Retriever and full-length SNX17 and reconstituted their association in the presence of a synthetic peptide corresponding to the LRP1 cytoplasmic tail containing the ^4470^NPTY^4473^ ØxNxx[Y/F] motif. SNX17 associated with Retriever coated beads, and this association was enhanced in a dose-dependent manner by inclusion of the LRP1 peptide (Fig. 4G, Supp. Fig. 2F). In control experiments, a corresponding LRP1 peptide carrying a Y4473A mutation in the ØxNxx[Y/F] motif failed to enhance SNX17 association to Retriever. Collectively, these data support the proposed model whereby an autoinhibited SNX17 conformation is released through a competitive interaction with ØxNxx[Y/F] cargo, which allows for the subsequent association of SNX17 and bound cargos with Retriever and the Commander super-assembly.

### SNX17 is not enriched with Commander at endosomal retrieval sub-domains

A number of studies have shown that specific endosomal membrane sub-domains are enriched for either degradative components such as ESCRT or recycling machinery including Retromer, Retriever, the CCC complex, and WASH-dependent actin patches (Puthenveedu et al., 2010; Temkin et al., 2011; Derivery et al., 2012; Gomez et al., 2012; van Weering et al., 2012; Bowman et al., 2016; Takatori et al., 2016; Varandas et al., 2016; McNally et al., 2017; Norris et al., 2017; Kvainickas et al., 2019; Singla et al., 2019; reviewed in Norris and Grant, 2020). We therefore explored the importance of SNX17 coupling to Retriever and cargo for their endosomal association and sub-domain organization. Like other cargo binding coat complexes, for example AP2 (Jackson et al., 2010), the endosomal localization of SNX17 requires avidity-based co-incident sensing of phosphoinositides, specifically phosphatidylinositol 3-monophosphate (PI(3)P) and ØxNxx[Y/F]-containing cargo proteins (Ghai et al., 2013). Consistent with this, the FERM-domain mutant GFP-SNX17(V380D) that exhibited decreased cargo-binding failed to localize to the endosomal network when transiently transfected into RPE1 cells (Fig. 4H). In contrast, the enhanced cargo-binding GFP-SNX17(F462A) mutant displayed an even more pronounced endosomal association, as evidenced by increased co-localisation with the endosomal marker EEA1 compared with wild-type GFP-SNX17 (Fig. 4H). These results suggest that SNX17 can cycle between the cytoplasm and the endosomal membrane through low affinity sensing of PI(3)P with the density of incoming ØxNxx[Y/F]-containing cargo providing an additional affinity to prolong the endosomal residency of SNX17 and release the autoinhibited conformation thereby facilitating coupling to Retriever and the Commander assembly.

To examine the endosomal organization of SNX17 and its relationship to Retriever association we used a VPS35L KO cell line engineered to stably express control GFP, or re-express GFP-VPS35L and GFP-VPS35L(R248A) – a mutant that retains Retriever assembly but inhibits binding to SNX17 (Fig. 2C) – uncoupling SNX17 mediated cargo recognition from the downstream process of cargo recycling. To evaluate the relative organization of GFP-VPS35L with endogenous SNX17 and other endosomal markers, we analyzed multiple endosomes through high-resolution confocal immunofluorescence imaging, and plotted the normalized average fluorescence intensity profiles to evaluate the relative distributions of endosomal markers within a single endosome. Endogenous SNX17 displayed a general distribution over the bulk of EEA1-positive endosomal membranes (Fig. 5A). In contrast GFP-VPS35L and GFP-VPS35L(R248A) were enriched on one or more foci of the SNX17-labelled endosomes, and co-localized with endogenous COMMD1 (Fig. 5B) and FAM21 (Supp. Fig. 3A), markers of the CCC and WASH complexes respectively. Retriever, CCC and WASH complex markers also co-localized with the core Retromer component VPS35 and SNX1, an ESCPE-1 subunit that drives endosomal tubulation during the biogenesis of transport carriers (van Weering et al., 2012, Simonetti et al., 2019; Lopez-Robles et al., 2023) (Supp. Fig. 3B, C). These foci therefore represent the previously described retrieval sub-domains from where cargo-enriched transport carriers exit the endosome for transport to their destination (Puthenveedu et al., 2010; Temkin et al., 2011; Bowman et al., 2016; Varandas et al., 2016; McNally et al., 2017). Our data shows that the association of SNX17 (and cargo) with Retriever was not a pre-requisite for sub-domain organization, as GFP-VPS35L(R248A) retained localization to the retrieval sub-domain. We also noticed that VPS35L KO cells displayed a partial perturbation in the sub-domain organization of FAM21, SNX1 and VPS35 (Supp. Fig. 3A-C). Interestingly, in VPS35L KO cells, these markers occupied multiple, less well-defined foci on the EEA1 and SNX17 endosomal membrane with greater localization overlap with EEA1 or SNX17 (Supp. Fig. 3A-C). Altogether these data indicate: (i) that VPS35L and the Retriever complex plays a role in organizing endosomal retrieval sub-domains; (ii) that cargo sensing by SNX17 can promote its recruitment to EEA1-positive endosomes but is not a pre-requisite for the formation of the retrieval sub-domain; and (iii) whilst present SNX17 is not enriched in these retrieval sub-domains. Rather it appears that the transient coupling of cargo bound SNX17 to Retriever and Commander may serve to handover cargo into a pre-existing retrieval sub-domain for endosomal exit and recycling to the cell surface (Fig. 5C).

**Figure 5.**
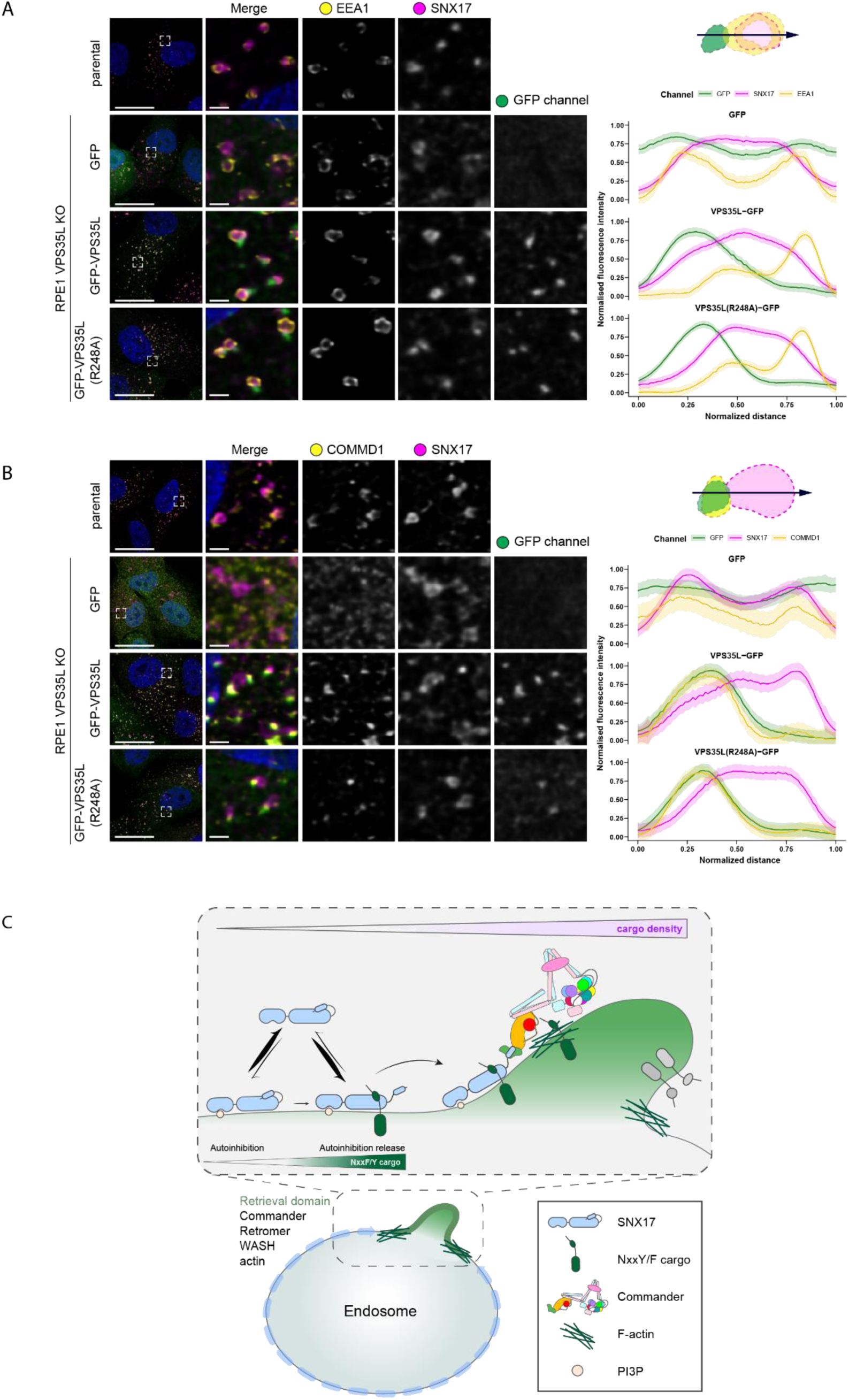
Retriever complex resides on retrieval subdomain of endosome and colocalises with markers of the CCC complex. **(A-B)** VPS35L KO RPE1 cells were lentiviral transduced with GFP, VPS35L-GFP or VPS35L-GFP(R248A). The localisation of GFP-tagged proteins was compared to the localisation of endogenous endosomal markers SNX17, EEA1 (A) and COMMD1 (B). Representative high-resolution confocal microscopy images are shown. The relative distributions of endosomal markers were evaluated in ImageJ by generating fluorescence intensity line profiles. Line profiles of 30 endosomes from 3 independent experiments were analysed in Rstudio, where the lengths of line scans and raw fluorescence intensities were normalised and averaged. The average profiles are shown on the right. The shading corresponds to the 95% confidence interval. **(C)** Model of SNX17-Commander association and its regulation through ØxNxxY/F cargo-density sensing and endosomal subdomain organisation. SNX17 associates with the endosomal membrane that is enriched for PI3P. We hypothesize that endosomal localization is enhanced through the binding of transmembrane cargoes. With increasing density of ØxNxxY/F cargo, auto-inhibitory sequence within SNX17 tail is displaced, consequently enabling the presentation of SNX17 carboxy-tail to the conserved VPS26C:VPS35L interface. The direct binding of cargo-bound SNX17 to Retriever ultimately leads to Commander-WASH complex-mediated recycling of cargo back to the plasma membrane. For simplicity, other endosomal sorting complexes, such as WASH, ESCPE-1 and Retromer are not shown.

In summary, by identifying the molecular details of SNX17 coupling to Retriever we have revealed the evolutionarily conserved mechanism through which hundreds of integral membrane proteins, including integrins and lipoprotein receptors, enter the Commander retrieval and recycling pathway. Importantly, we establish that cargo binding facilitates SNX17 endosomal association, and that cargo occupancy relieves an autoinhibited SNX17 conformation to promote Retriever association and the entry of cargo into a pre-existing retrieval sub-domain for the promotion of Commander mediated cell surface recycling. Overall, our study provides fundamental mechanistic and regulatory insight into the role of the SNX17-dependent Commander retrieval and recycling pathway during essential cellular processes ranging from directed cell migration through to cholesterol homeostasis.

## LIMITATIONS OF THIS STUDY

Our work provides the mechanism for SNX17 coupling to Retriever and the Commander super-assembly, the influence of cargo occupancy on this association and the importance for entry into the retrieval sub-domain. However, while we have provided strong biochemical and functional data supportive of the proposed coupling mechanism full validation will require high resolution structural analysis of the SNX17-Retriever and SNX17-Commander assemblies. Further experiments will also be required to broaden our mechanistic understanding of the dynamics of cargo handover into the retrieval sub-domain and the process of tubular-based exit, and how these events are controlled in response to changes in the cellular state.

## MATERIAL AND METHODS

### Cell culture

HEK293T and RPE1-hTERT cell lines were maintained in DMEM (Sigma, D5796) with 10% (v/v) fetal bovine serum (Sigma, F7526) in presence of penicillin/streptomycin (Gibco) at 37°C with 5% CO_2_.

### VPS35L CRISPR KO cell line generation

A RPE1-hTERT cell line lacking VPS35L was generated using CRISPR/Cas9 technology. The target sequence of CCTGTTTCTTGTTCGAGAGCTTC on exon28 (NM_020314.7) was inserted into the px458 plasmid (Addgene plasmid no. 48138) and transfected to cells using PEI. After 48 hrs of transfection, GFP-positive cells were sorted by FACS for single clone isolation. VPS35L-KO was confirmed by WB analysis.

### Site-directed mutagenesis

Primers for site-directed mutagenesis were designed using Agilent QuikChange Primer design tool. QuikChange II Site-Directed Mutagenesis Kit (Agilent, 200523-5) was used for mutagensis of GFP-VPS35L constructs following the manufacturer’s protocol. All other site-directed mutagenesis was carried out using Q5 High-Fidelity 2X Master Mix (NEB, M0492) following the manufacturer’s protocol. Nonmutated template DNA was digested after the PCR SDM reactions by 1h incubation with Dpn1 enzyme at 37°C. This DNA was used for bacterial transformation into XL10 Gold (Agilent, 200315) chemically competent cells according to manufacturer’s instructions. The bacteria were grown on appropriate antibiotic-containing agar plates. Full open-reading frames were sequenced to verify the results of mutagenesis.

### Gibson assembly

To subclone VPS35L-GFP into lentiviral pLVX vector, wild-type or mutant VPS35L sequences were amplified using Q5 High-Fidelity 2X Master Mix (NEB, M0492) following the manufacturer’s protocol. After amplification, PCR samples were resolved on agarose gel and amplified fragments extracted from gel using GFX PCR DNA and Gel Band purification kit (GE Healthcare, 28-9034-70). The purified DNA or 1µg of plasmid DNA were digested with XmaI and XbaI for 1 h at 37°C in 1x CutSmart buffer and nuclease-free water. To eliminate self-ligation, the plasmid was also treated with 1.5 µl of quick-CIP (NEB, M0525). Following digestion, DNA was again purified as above and a ligation reaction between 1:6 ratio of backbone:vector was carried out with T4 DNA ligase (Invitrogen, 15224017). Ligation mixture was incubated at 10, 20 and 30°C for 30s, for a total of 200 cycles.

### Recombinant protein expression and purification

Retriever with a VPS29-His tag and a Strep-VPS26C or Halo-VPS26C tag was expressed in Sf21 insect cells and purified as described previously (Healy et al., 2023). A gene encoding full-length human SNX17 with an N-terminal Strep-tag and HaloTag® was codon optimised for S. frugiperda and purchased from Twist Biosciences (San Francisco, CA). The gene was cloned into pACEBac1 using BamHI/HindIII restriction sites and the resulting pACEBac1-Strep-Halo-SNX17 vector was used to generate V1 and V2 baculoviruses as described previously (Healy et al., 2023). For protein expression, 4mL V1 or V2 virus was added to Sf21 cells at a density of 0.5-1.0x106mL in 400mL Sf-900 II SFM (Thermo). Cells were harvested 72 hours post-infection and cell pellets were stored at -80°C until use. For SNX17 purification, insect cells were resuspended in cold lysis buffer (50mM HEPES pH7.2, 150mM NaCl, 2mM β-mercaptoethanol, 0.1% (v/v) Triton X-100, EDTA-free protease inhibitor (Pierce)) and lysed on ice using a 130-Watt Ultrasonic Processor (Cole-Palmer) for 2 mins 30s using a 10s-on 30s-off cycle. Insoluble material was removed by centrifugation at 20000rpm in a JA-20 fixed-angle rotor (Beckman Coulter) for 25 mins at 4°C, then the soluble lysate was incubated with 1mL equilibrated Strep-Tactin® Sepharose® resin (IBA Lifesciences) rotating for 1 hour at 4°C. After binding, the resin was washed 3x in lysis buffer and protein was eluted with a further 3 washes in elution buffer (50mM HEPES pH7.2, 150mM NaCl, 2mM β-mercaptoethanol, EDTA-free protease inhibitor (Pierce), 2.5mM desthiobiotin (IBA Lifesciences)). Eluates containing protein were concentrated to <500µL and size-exclusion was performed at 4°C using an ÄKTA Pure Protein Purification System (GE healthcare) and Superose® 6 Increase 100/300 GL column (Cytiva), with 0.5mL fractions collected using an F9-C Fraction Collector (Cytiva). Fractions containing SNX17 were pooled and concentrated.

SNX17 full length (residue 1-470) and SNX17ΔC (residue 1-390) were synthesised with an intramolecular decaHis tag between residue 335 and 346 by GeneUniversal and subcloned into pET28a (XbaI and XhoI). GST-LRP1 was available in a pgex4T-2 vector, GST was produced from expression of a native pGEX6P-1 vector (Ghai et al., 2013). These plasmids were transformed into *E.coli* BL21 DE3 competent cell (New England Biolabs) and plated on agar plates containing Ampicillin or Kanamycin. Clones from this agar plate were grown overnight in 30 mL of LB broth. 5 mL from these cultures was added to 1 L of LB supplemented with 40 mM NH_4_Cl, 4 mM MgCl_2_, 4 mM NaSO_4_, 2.5% glycerol and 30 mM α-lactose and grown at 25°C for 24 h. Cells were harvested by centrifugation at 6000 x g for 10 min at 4°C and the harvested cell pellet was resuspened in 50 mM Tris pH 8.0, 500 mM NaCl, 5 mM imidazole, 2 mM β-mercaptoethanol, 10% glycerol, 50 μg/mL benzamidine and 100 units of DNAseI. Cells were lysed by cell disruption at 35 kPSI and clarified by centrifugation 50,000 x g for 30 mins at 4°C. Talon or glutathione Sepharose (Clonetech) was used to isolated SNX17 constructs and GST constructs, respectively. His tagged constructs were eluted via 500 mM imidazole and GST-tagged proteins were removes via 50 mM glutathione, other buffer components were as above. These eluted proteins were subsequently passed through a superdex s75 16/60 column attached to an AKTA Pure system (GE healthcare) in 50 mM Tris pH 8.0 and 300 mM NaCl.

### *In vitro* SNX17-Retriever interaction assays

0.1 mg/mL purified Retriever with a VPS29-His tag was mixed with 0.075 mg/mL purified SNX17 (WT) or SNX17 (L470G) in a total volume of 0.1 mL cold lysis buffer (50 mM HEPES pH7.2, 150 mM NaCl, 2 mM β-mercaptoethanol, 0.1% (v/v) Triton X-100, EDTA-free protease inhibitor (Pierce)). LRP1 (NPXY) (Biotin-GRMTNGAMNVEIGNPTYKMYEGGEPDDG) and LRP1 (NPXA) (Biotin-GRMTNGAMNVEIGNPTAKMYEGGEPDDG) peptides corresponding to human LRP1 amino acid residues 4458-4483 were purchased from GenScript and, where indicated, added to a final concentration of 3 µM or 30 µM. Protein mixtures were incubated with 20 µL equilibrated TALON® Superflow beads (Cytiva) rotating for 1 hour at 4°C, then beads were washed 3x in cold wash buffer (50 mM HEPES pH7.2, 150 mM NaCl, 2 mM β-mercaptoethanol, 0.1% (v/v) Triton X-100, 10 mM imidazole, EDTA-free protease inhibitor (Pierce)). Washed beads were resuspended in 4x SDS loading dye + 2.5% β-mercaptoethanol for analysis by SDS PAGE followed by Coomassie staining and western blotting.

In the presence of 20 μL of glutathione Sepharose (Clonetech) 2 nmol SNX17 or SNX17ΔC was mixed with 1 nmol GST-LRP1 for 5, 10, 30 or 60 mins and 1 nmol GST for 60 mins. At each time point the protein resin mixture was centrifuged at 5000 x g for 1 min and the supernatant was removed to stop further interaction. After 60 min glutathione resin was washed 5 times with a buffer containing 20 mM Tris pH 8.0, 300 mM NaCl, 2 mM β-mercaptoethanol, 10% glycerol, 0.5% igepal and EDTA-free protease inhibitor (Pierce). All buffer was removed, and beads were resuspended in 30 μL of buffer (100 NaCl, 20 mM Tris pH 8.0) and 10 μL 4x SDS loading dye before incubation at 100°C for 5 min. 10 μL of sample was loaded onto precast SDS-PAGE gels (Novex) and after running for 43 min at 165v were stained using Coomassie blue (Sigma).

### Transient transfection

For GFP trap experiments and lentiviral particle production, HEK293T cells were transiently transfected using PEI (polyethyleneimine). 10 ml of Opti-MEM were equally split between 2 sterile tubes. The first tube was used to dilute 15 µg plasmid DNA (or as described in ‘Lentiviral particle generation’), and the second to dilute PEI at 3:1 PEI:DNA ratio. The PEI dilution was sterilised through 0.2 µm filter. The total contents of both tubes were then mixed and incubated for 15 min prior to tranfection of cells. The cells were incubated with DNA:PEI mixture in Opti-MEM for 6h, and after incubation, this was replaced for normal growth media for 24 or 48h. For imaging experiments, RPE1-hTERT cells were transfected using Lipofectamine LTX Reagent with PLUS Reagent (Invitrogen, A12621) according to manufacturer’s instructions. Briefly, 200 µl of Opti-MEM were split equally between 2 sterile tubes. The first tube was used to dilute 0.5 µg plasmid DNA in the presence of 3 µl Plus reagent. The other tube was used to dilute 3 µl of Lipofecatmine. The total contents of both tubes were then mixed and incubated for 5 min at room temperature. Following the incubation, transfection mixture was added drop wise to cells grown on coverslips in 6-well plates and cells were grown for 24h before fixation.

### Lentiviral particle and stable cell line generation

To generate lentivirus, HEK293T cells were grown in 15 cm dishes and transfected with 15 μg of PAX2, 5 μg pMD2.G and 20 μg of lentiviral expression vector using PEI transfection. After the 48h incubation, the growth media containing the lentivirus was harvested and filtered through a 0.45 μm filter. 6-wells of VPS35L KO RPE1-hTERT cells were transduced at 25% confluency with varying volumes of lentiviral media. After 48h, cells were treated with 15 μg/ml of puromycin to select cells with successfully integrated lentiviral constructs.

### Quantitative GFP-nanotrap

Transiently transfected HEK293T cells, expressing GFP or GFP-tagged proteins were rinsed twice with ice-cold PBS and then lysed in a buffer containing 0.5 % NP-40, 50 mM Tris pH7.5 in ddH_2_O (in GFP-SNX17 pull-downs) or PBS (co-transfections). Supernatant was collected after 10 min centrifugation at 15000 rpm. 30 μl of supernatant was removed to serve as whole-cell input and diluted in 1:1 ratio with 4x loading buffer containing 2.5 % β-mercaptoethanol. The remaining supernatant was incubated with 25 μl of GFP-nanotrap (gta-20, Chromotek) beads for 1h at 4°C. The beads were then washed twice in a buffer containing 0.25 % NP-40, 50 mM Tris pH7.5 in PBS and once in 50 mM Tris pH7.5 in PBS (or diluted in ddH_2_O for GFP-SNX17 pull-downs). Between the washes, the beads were collected at the bottom of the tube by 1min centrifugation at 2000 rpm. Beads were resuspended in a 2x loading buffer, containing 2.5 % β-mercaptoethanol and all samples were denatured at 95°C for 10 minutes.

### Cell surface protein biotinylation

RPE1-hTERT cells were washed generously with ice-cold PBS prior to labelling with cell-impermeable 0.2 mg/mL Sulfo-NHS-SS Biotin (ThermoFisher Scientific, no. 21217) in PBS (pH 7.4). The biotinylation reaction was then quenched by incubating the cells in TBS for 10 minutes, During the labelling, quenching and washing steps, cells were incubated on ice to prevent endocytosis and unspecific labelling of intra-cellular proteins. After quenching, cells were lysed in lysis buffer containing 2% triton x-100 with protease inhibitors in PBS. Supernatant was collected after 10min centrifugation at 15000 rpm, and total protein levels analysed using BCA reaction. Inputs were collected and stored separately. Equal amounts of protein were incubated with Streptavidin beads (GE Healthcare, USA) for 30 min at 4°C. The samples were then washed once in PBS with 1% triton x-100, twice in PBS with 1% triton x-100 with 1M NaCl and once with PBS. Between the washes, the beads were collected at the bottom of the tube by 1min centrifugation at 2000 rpm. Beads were resuspended in a 2x loading buffer, containing 2.5 % β-mercaptoethanol and all samples were denatured at 95°C for 10 minutes.

### Immunoblot

Equal amounts of samples were loaded onto NuPAGE® 4-12% gradient Bris-Tris gels (NP0322BOX, Invitrogen) and resolved at 130V. Proteins were transferred onto a methanol-activated PVDF membrane at 100V for 75 minutes in a transfer buffer containing 25 mM tris, 192 mM glycine, and 10% methanol. After transfer, membranes were blocked in 10% milk in TBST (tris-buffered saline with Tween: 150 mM NaCl, 10 mM tris pH7.5, 0.1% Tween-20) for 1 hour at room temperature. They were then washed in TBST and incubated overnight in primary antibody diluted in 3% BSA in TBST. After the incubation, the membrane was washed three times in TBST for 5 minutes and incubated with fluorophore-conjugated secondary antibodies diluted 1:20000 in 5% milk for 1 hour at room temperature. Prior to imaging on LI-COR Odyssey CLx system, membranes were washed generously in TBST buffer. Quantification of band intensities was performed in Image Studio Lite software and GraphPad Prism 8 was used for the statistical analysis.

### Microscopy sample preparation

Cells, grown on 13mm glass coverslips, were fixed with 4% PFA (Invitrogen, 28906) in PBS for 15min at room temperature. To remove fixative, cells were washed three times before permeabilisation in 0.1% (v/v) Triton X-100 in PBS or, in case of LAMP1 co-staining, 0.1% (w/v) saponin. Coverslips were then washed again three times with PBS before 20 min incubation with 1% (w/v) BSA in PBS. Coverslips were then incubated with primary antibodies diluted in 0.1 % (w/v) BSA in (with added 0.01% (w/v) saponin in case of saponin permeabilisation) for 1h at room temperature. After primary antibody detection, coverslips were washed generously in PBS, before secondary detection with Alexa Fluor-conjugated secondary antibodies and DAPI for 1 h at room temperature. The coverslips were finally washed three times with PBS and once in destillled water before mounting onto glass microscopy slides using Fluoromount-G (Invitrogen, 004958-02) and kept refrigerated before analysis on the microscope.

### Confocal microscopy and image analysis

Fixed cells were imaged on Leica SP8 multi-laser point scanning confocal microscope, using a 63x NA1.4 UV oil-immersion lens. Endosomal sub-domains were resolved using Leica ‘Lightning’ mode for adaptive deconvolution to improve lateral resolution. Leica LAS X software was used for the acquisition of images. Pearson’s colocalisation coefficients were determined using Volocity 6.3.1 software (PerkinElmer) with automatic Costes background thresholding. Representative images for colocalisation images were also prepred using the Volocity software. For line fluorescence analysis, high-resolution microscopy images were opened in Fiji software. 30 endosomes from 3 independent experiments per condition were analysed. Briefly, a line was drawn from ‘retrieval sub-domain’ to the end of ‘endosomal core’ and fluorescence intensity along line for each channel obtained using ‘Plot Profile’ function. In R studio software, the fluorescence intensities were normalies to maximum fluorescence intensity of respective channel and endosome. The data was analysed and represented in R studio software using ‘ggplot2’ package. Representative images for subdomain organisation were prepared in Fiji.

### AlphaFold2 modelling

All AlphaFold2 models were generated using AlphaFold multimer version 3 (Jumper, 2021 and Evans 2022) implemented in the ColabFold interface available of the Google Colab platform (Mirdita, 2022). The models present in this study include: Retriever (VPS35L, VPS26C and VPS29) in complex with SNX17^400-470^ (Model archive: ), SNX17^1-470^ in complex with LRP1^4460-4490^ (Model archive: ), Retriever in complex with ^SNX171-470^ (Model archive: ), *D.melanogaster* Retriever in complex with SNX17^1-470^ (Model archive: ) and SNX17^1-470^. 5 independent models were generated for each complex and the quality of the predicted complexes was assessed through examination of iPTM score and the predicted alignment error plot.

### Antibodies

| Target | Supplier | Cat. Number |
| --- | --- | --- |
| CCDC22 | Proteintech | 16636 |
| CCDC93 | LSBio | C336997 |
| COMMD1 | Sigma Aldrich | WHO150684M1 |
| EEA1 | BD Biosciences | 610457 |
| FAM21 | Gift from Dan Billadeau | N/A |
| GFP | Roche | 11814460001 |
| Itgα5 | Abcam | ab150361 |
| Itgβ1 | BD Biosciences | 610467 |
| LAMP1 | Developmental Studies<br>Hybridoma Bank | H4A3 |
| LRP1 | Abcam | ab92544 |
| mCherry | Antibodies.com | A85306 |
| N-cadherin | Cell Signalling Technology | (13A9) 14215 |
| SNX1 | BD biosciences | 611482 |
| SNX17 (IF) | Sigma | HPA043867 |
| SNX17 (WB) | Proteintech | 10275 |
| VPS26C | Millipore/Sigma Aldrich | ABN87 |
| VPS29 | Santa Cruz | SC-398874 |
| VPS35 | Antibodies.com | A83699 |
| VPS35L | Abcam | ab97889 |
| β-actin | Sigma | A1978 |
| **Goat anti-Mouse IgG (H+L)**<br>**Cross-Adsorbed Secondary**<br>**Antibody, Alexa Fluor 680** | Invitrogen | A-21057 |
| **Goat anti-Rabbit IgG (H+L)**<br>**Secondary Antibody,**<br>**DyLight 800** | Invitrogen | SA5-35571 |
| **Alexa Fluor 568 anti-mouse**<br>**IgG** | Invitrogen | A10037 |
| **Alexa Fluor 568 anti-rabbit**<br>**IgG** | Invitrogen | A10042 |
| **Alexa Fluor 647 anti-mouse**<br>**IgG** | Invitrogen | A31571 |
| **Alexa Fluor 647 anti-rabbit**<br>**IgG** | Invitrogen | A31573 |
| **Alexa Fluor 647 anti-goat**<br>**IgG** | Invitrogen | A21447 |

### Primers

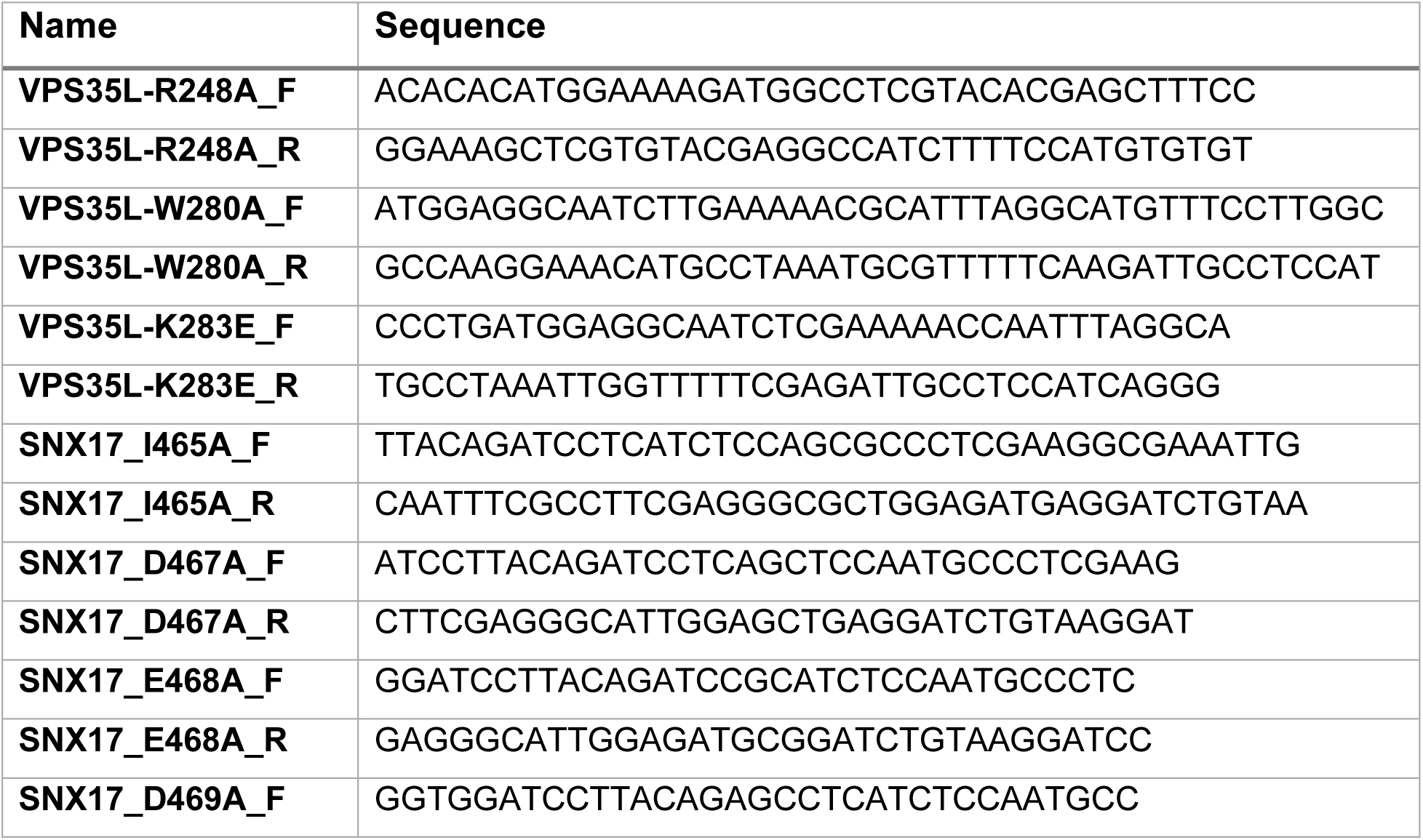
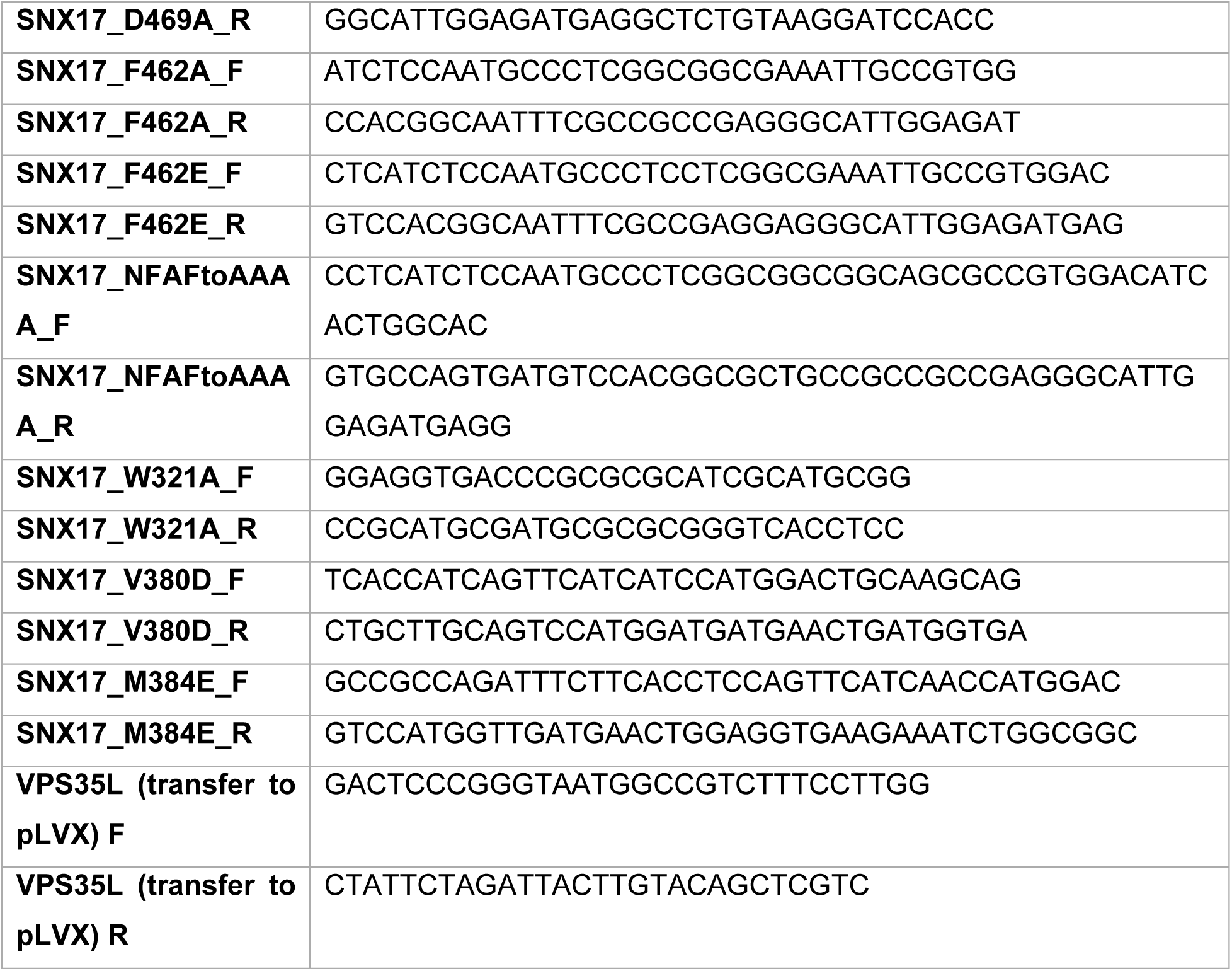

## ACKNOWLEDGEMENTS

We thank the Wolfson Bioimaging Facility at the University of Bristol for their support and Manu Derivery (MRC-LMB) for insightful discussions. Work in the Cullen laboratory is supported by the Wellcome Trust (104568/Z/14/Z and 220260/Z/20/Z), the Medical Research Council (MR/L007363/1 and MR/P018807/1), the Lister Institute of Preventive Medicine, and the award of a Royal Society Noreen Murray Research Professorship to P.J.C. (RSRP/R1/211004). R.B. is supported by the EndoConnect European Research Training Network (No. 953489). B.M.C. is supported by an Investigator Grant, Senior Research Fellowship and Project Grant from the National Health and MRC (APP2016410, APP1136021 and APP1156493). M.D.H is supported by a Dementia Australia Research Foundation (DARF) project grant.

## AUTHOR CONTRIBUTIONS

Cell-based biochemistry and analysis: R.B., M.D.H Protein purification and recombinant reconstitution: A.P.W., M.D.H., M.L. AlphaFold2 modelling: R.B., M.D.H. and B.M.C. Manuscript Writing - 1^st^ draft: R.B., B.M.C. and P.J.C; Final Version: all authors. Initial Concept: R.B., M.D.H., K.E.M., B.M.C. and P.J.C. Concept Development: all authors. Funding and Supervision: M.D.H., K.E.M., B.M.C. and P.J.C.

---

**Figure S1.**
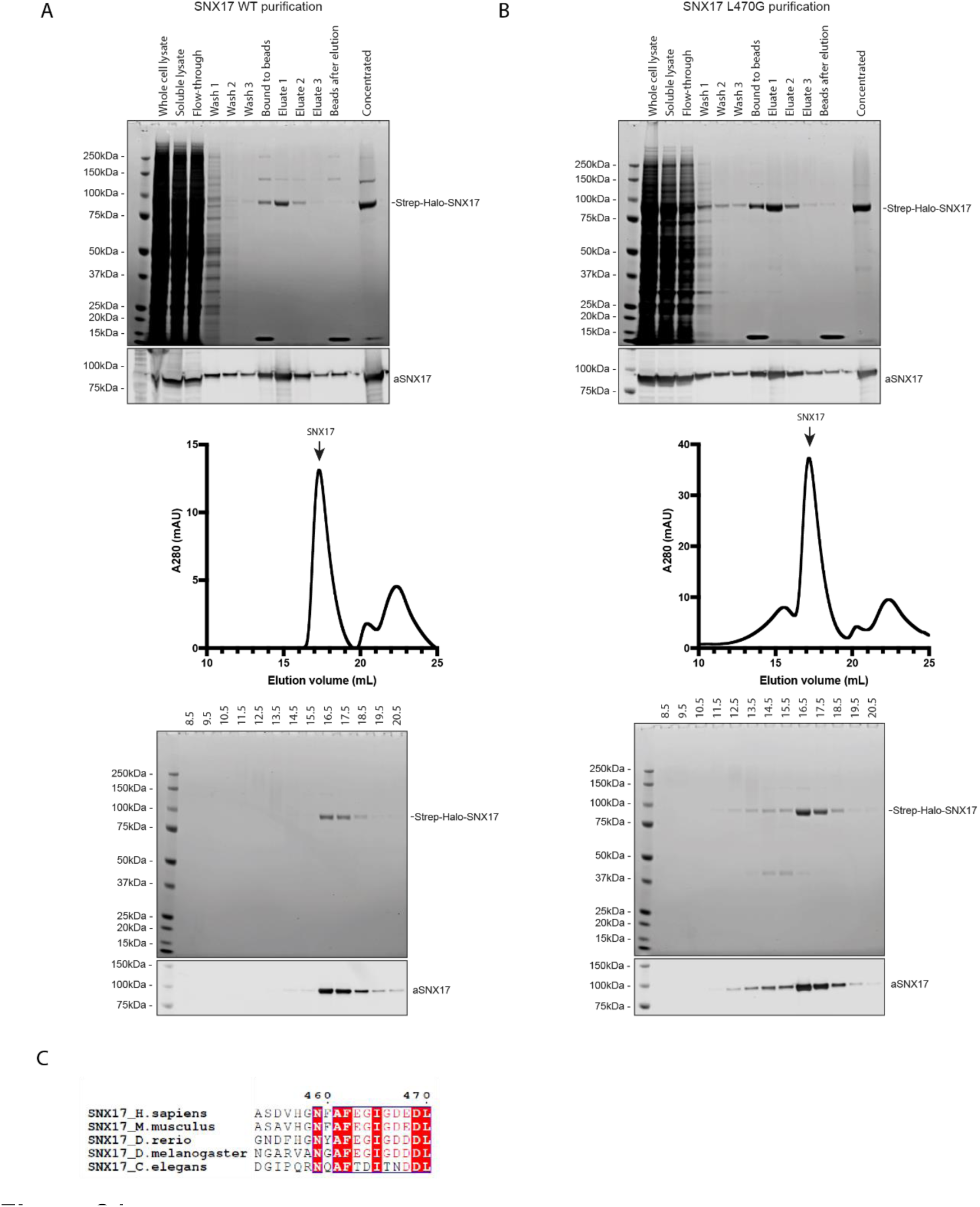
Purification of wild-type SNX17 **(A)** and SNX17(L470G) **(B)**. **(C)** Evolutionary conservation of C-terminus of SNX17 was evaluated by aligning full length SNX17 protein sequences from different model organisms in Clustal Omega online tool and depicted using ESPript3.0. Last 18 residues from each species are shown. The residue numbering corresponds to human SNX17 protein.

**Figure S2.**
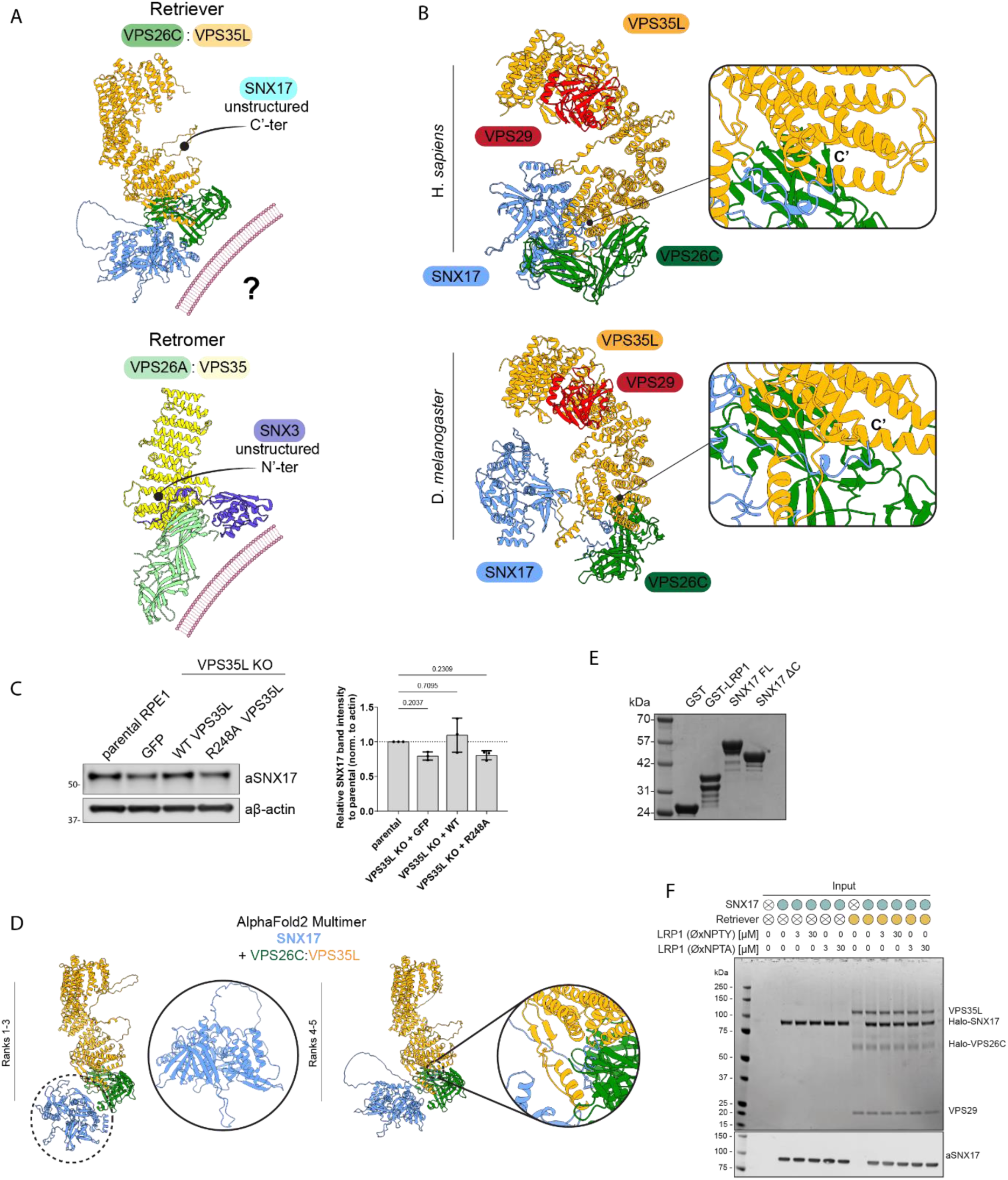
**(A)** Comparison of SNX17 binding to the VPS35L-VPS26C interface (AlphaFold2) and SNX3 binding to VPS35-VPS26A interface of the Retromer complex (X-ray, PDB 5F0J). Previously characterised membrane binding interface within retromer is depicted with the cartoon of phospholipid bilayer. The possible orientation of SNX17-Retriever complex relative to membrane is shown. **(B)** Comparison of AlphaFold2-predicted conformations of SNX17-Retriever complex in human and Drosophila emphasizes evolutionary conservation of the assembly. **(C)** VPS35L KO RPE1 cells were lentivirally transduced with GFP, VPS35L-GFP or VPS35L-GFP(R248A). Protein lysates were then resolved using immunoblot and whole-cell levels of SNX17 were compared in all samples. The quantification from 3 independent experiments is shown on the right. n = 3, 1-way ANOVA with Dunnett’s multiple comparison test, error bars represent s.d. **(D)** AlphaFold2 was used to predict the binding of full-length SNX17 to the VPS35L-VPS26C interface. Ranks were automatically assigned to the 5 predicted models, with ranks 1-3 predicting no binding between SNX17 and Retriever and showing unbound, monomeric SNX17, whereas ranks 4-5 predict the interaction between the unstructured C-terminus of SNX17 and VPS35L-VPS26C interface of the Retriever complex. **(E)** Input mixtures for Fig. 4G. **(F)** Input proteins for Fig. 4F.

**Figure S3.**
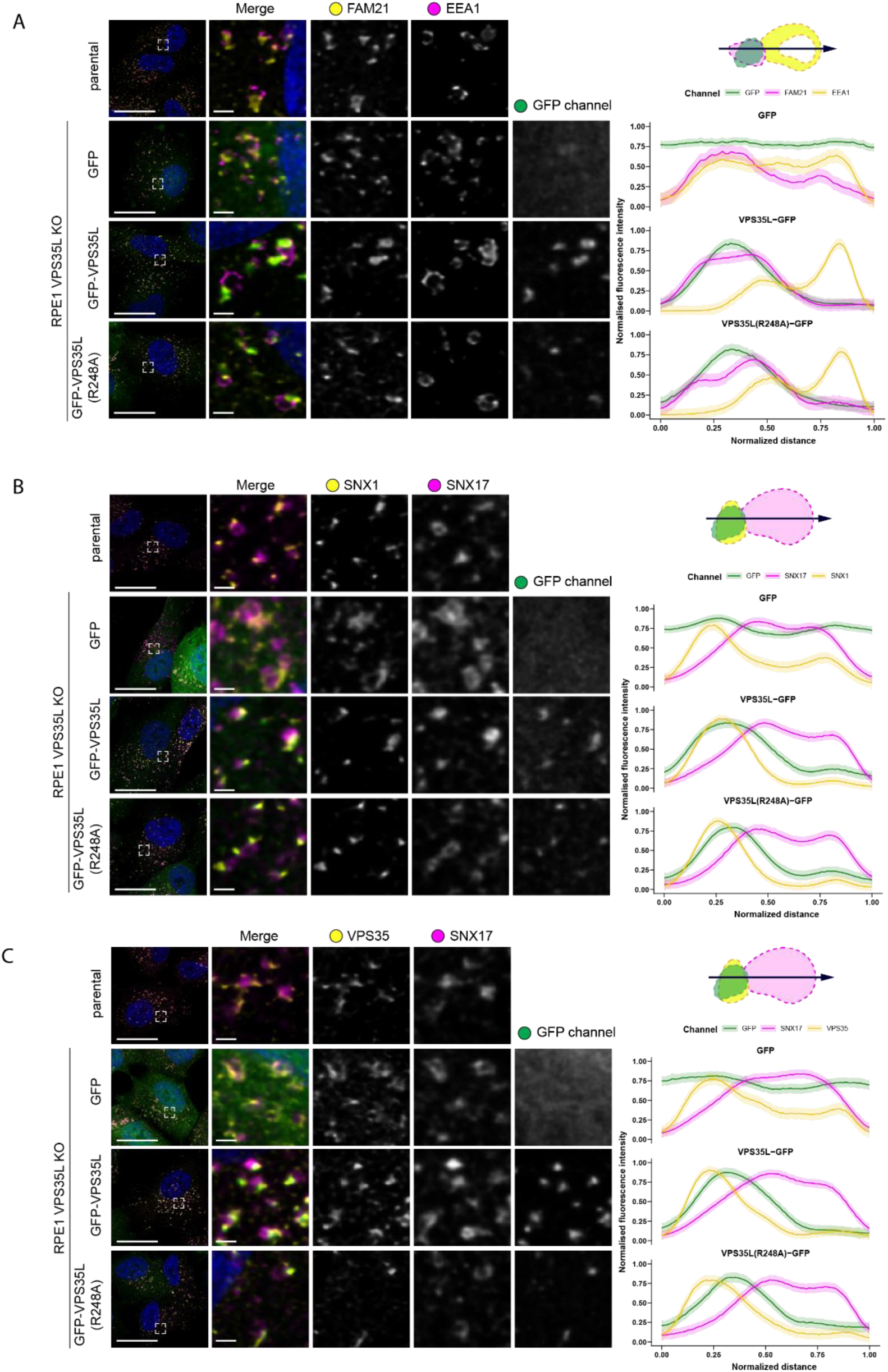
Retriever complex colocalises with the markers of the WASH, ESCPE and Retromer complexes. **(A-C)** VPS35L KO RPE1 cells were lentivirally transduced with GFP, VPS35L-GFP or VPS35L-GFP(R248A). The localisation of GFP or GFP-tagged proteins was compared to the localisation of endogenous endosome markers SNX17 and FAM21 (WASH complex subunit) (A), SNX1 (ESCPE-1 subunit) (B) or VPS35 (Retromer) (C). Representative high-resolution confocal microscopy images are shown. The relative distributions of endosomal markers were evaluated in ImageJ by generating fluorescence intensity line profiles. Line profiles of 30 endosomes from 3 independent experiments were analysed in Rstudio, where the lengths of line scans and raw fluorescence intensities were normalised and averaged. The average profiles are shown on the right. The shading corresponds to the 95% confidence interval. Scale bars shown for full image or inset correspond to 20 µm and 2 µm, respectively.

**Figure S4.**
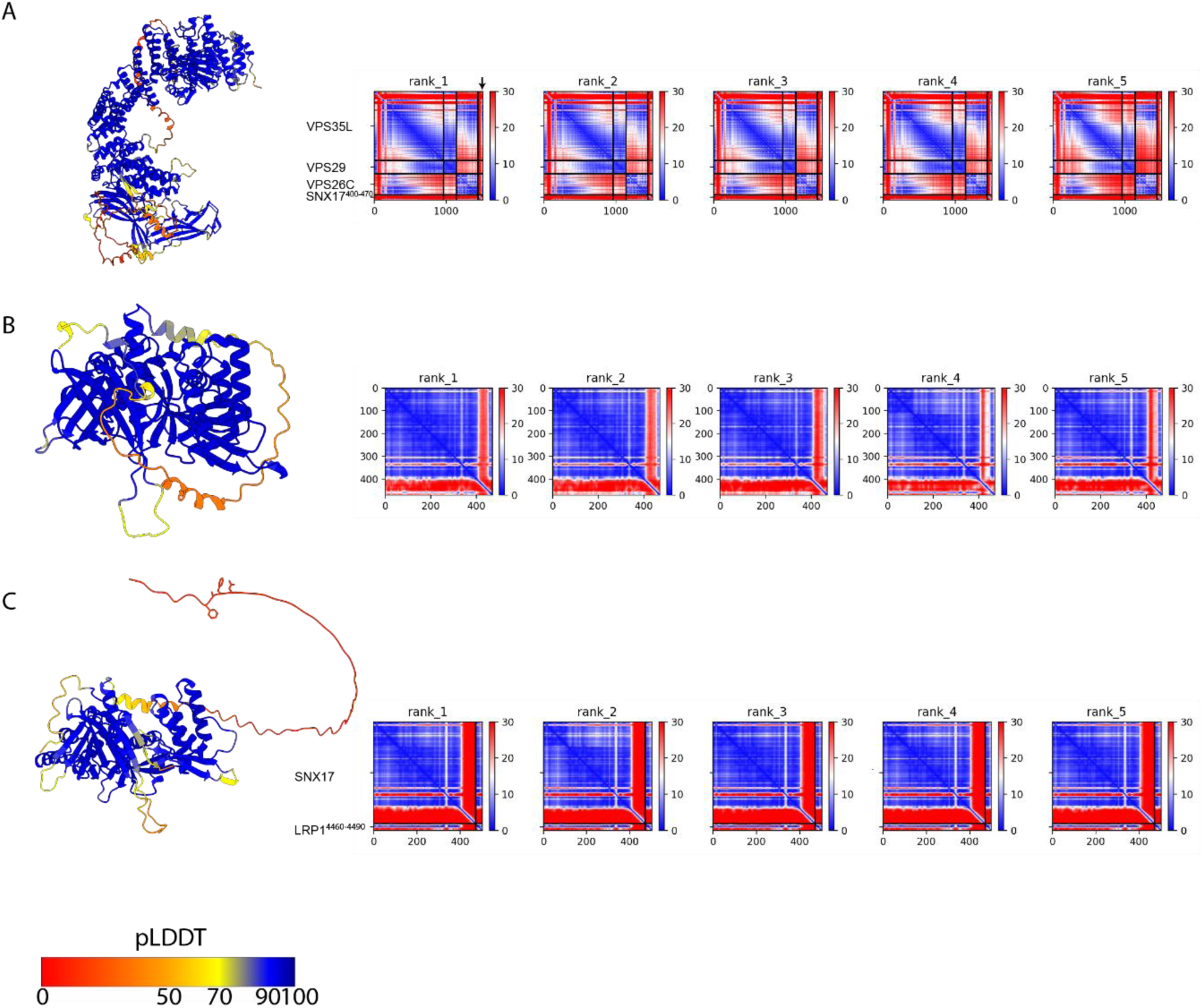
AlphaFold2 models and associated predicted alignment error plots. **(A)** AlphaFold2 prediction of Retriever (VPS35L, VPS26C, VPS29) and the residue 400 to 470 of SNX17, **(B)** SNX17 full length and **(C)** SNX17 full length modelled against the LRP1 carboxy-tail. Each structure model is coloured by pLDDT score, a measure of confidence where blue is a more confident prediction. Each structure is also accompanied by the predicted alignment error (PAE) plot for each of the five models. PAE plots show the correlation between any given residue in angstroms (Å). Blue indicates that two residues are highly correlated while red indicates no correlation.

## SOURCE DATA

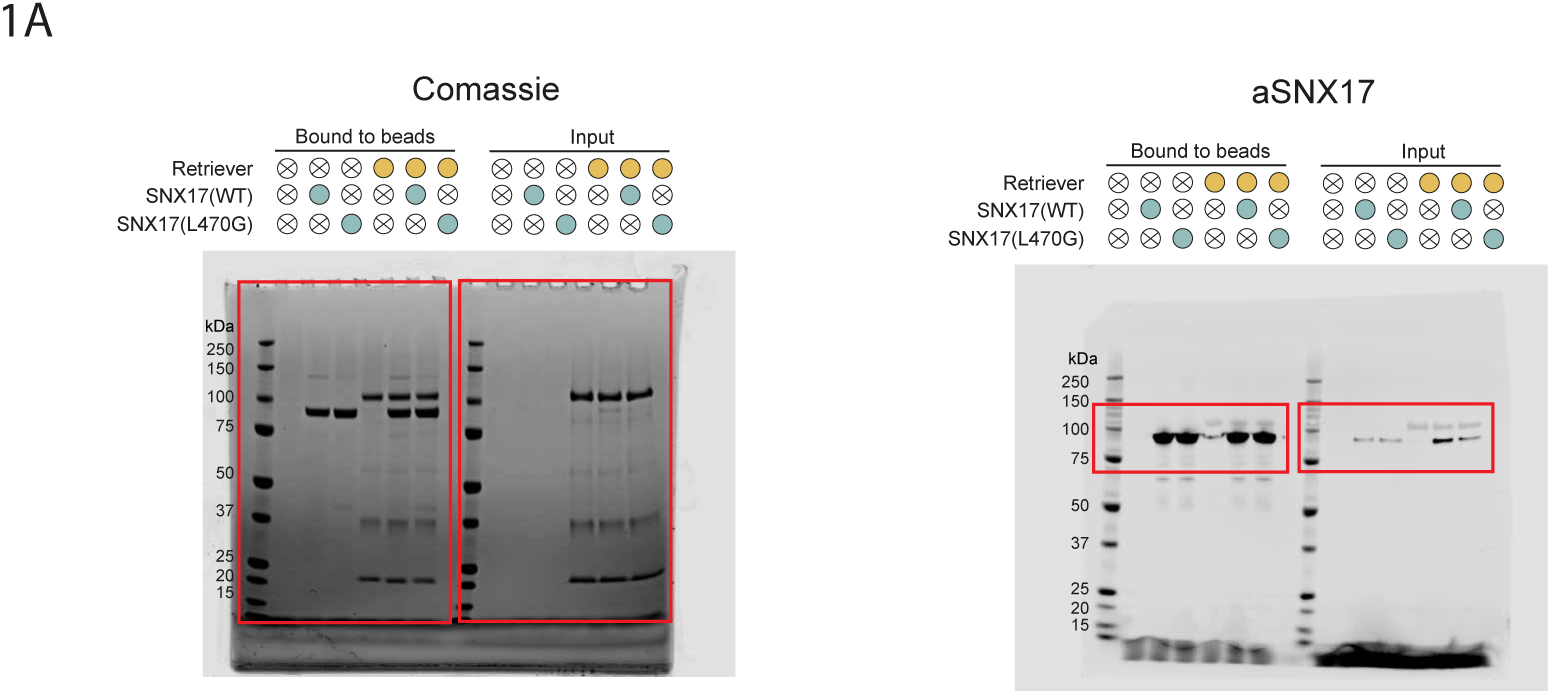

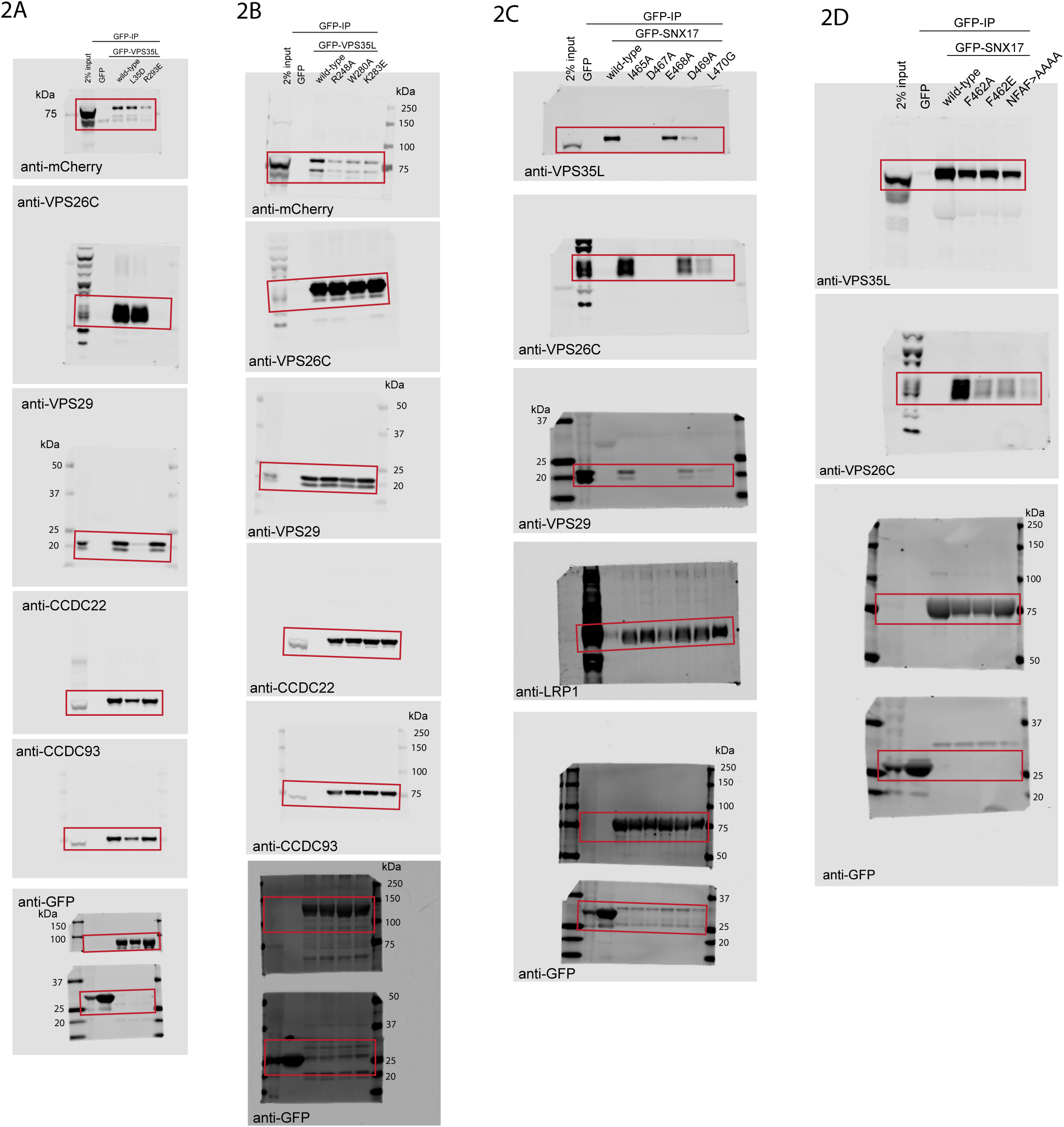

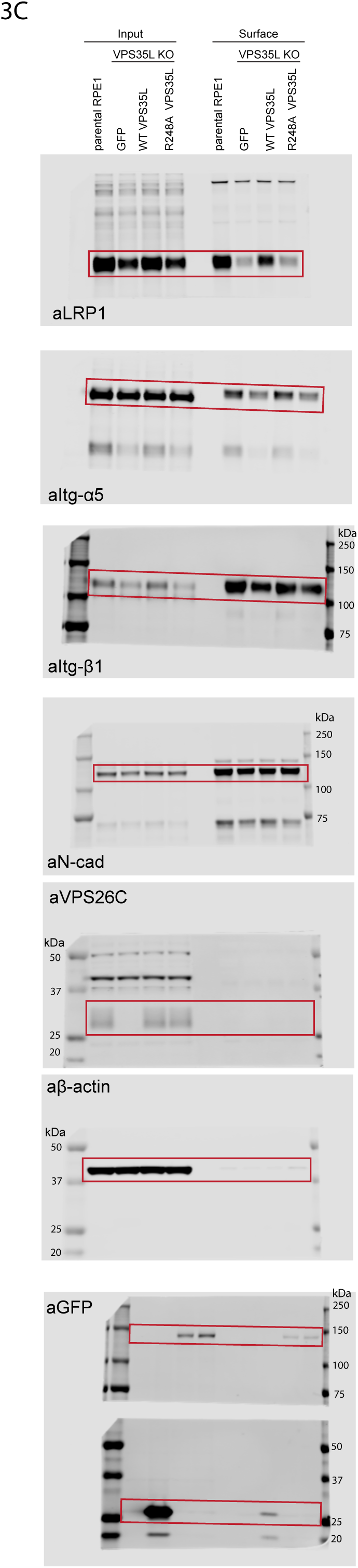

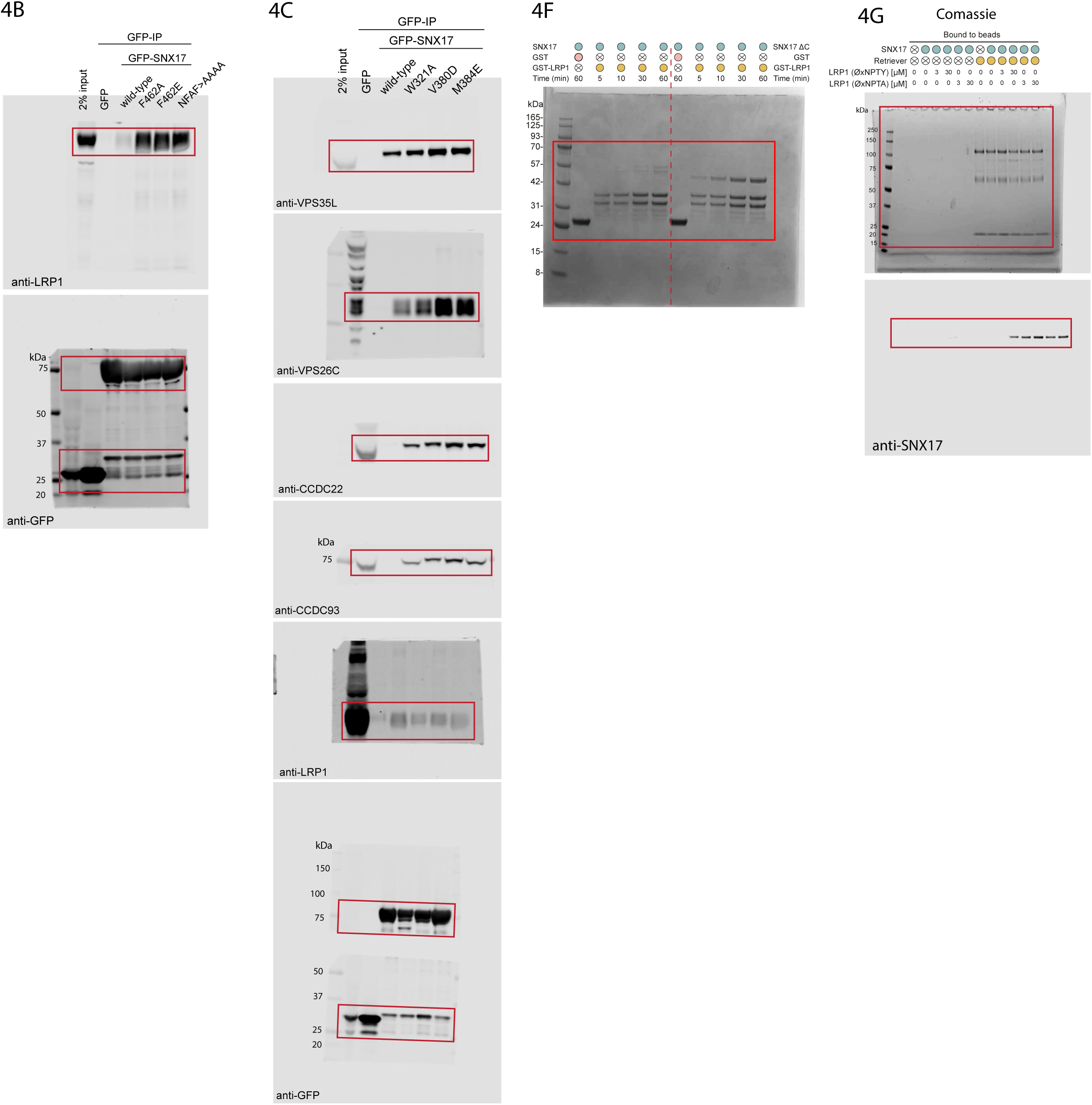

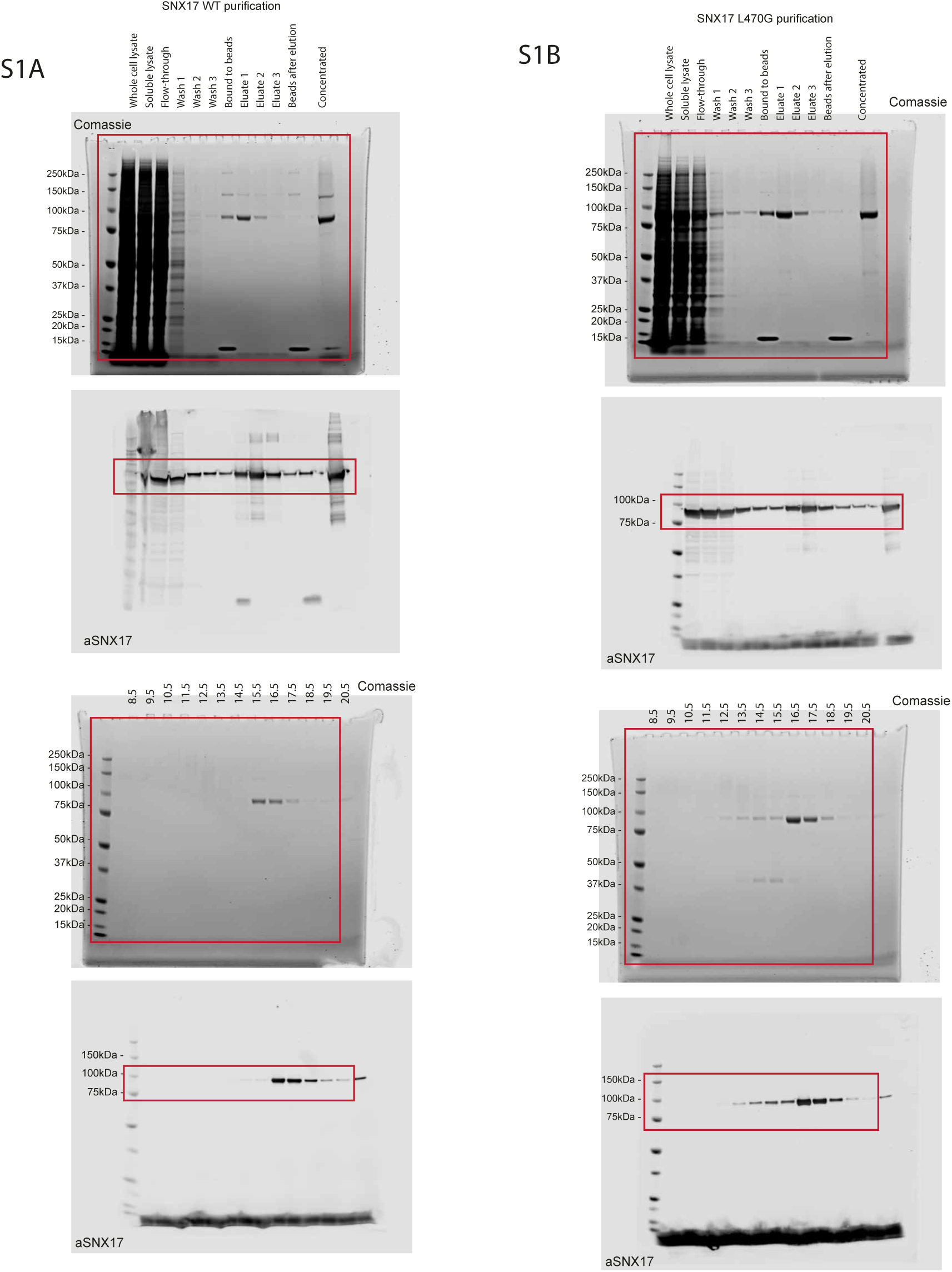

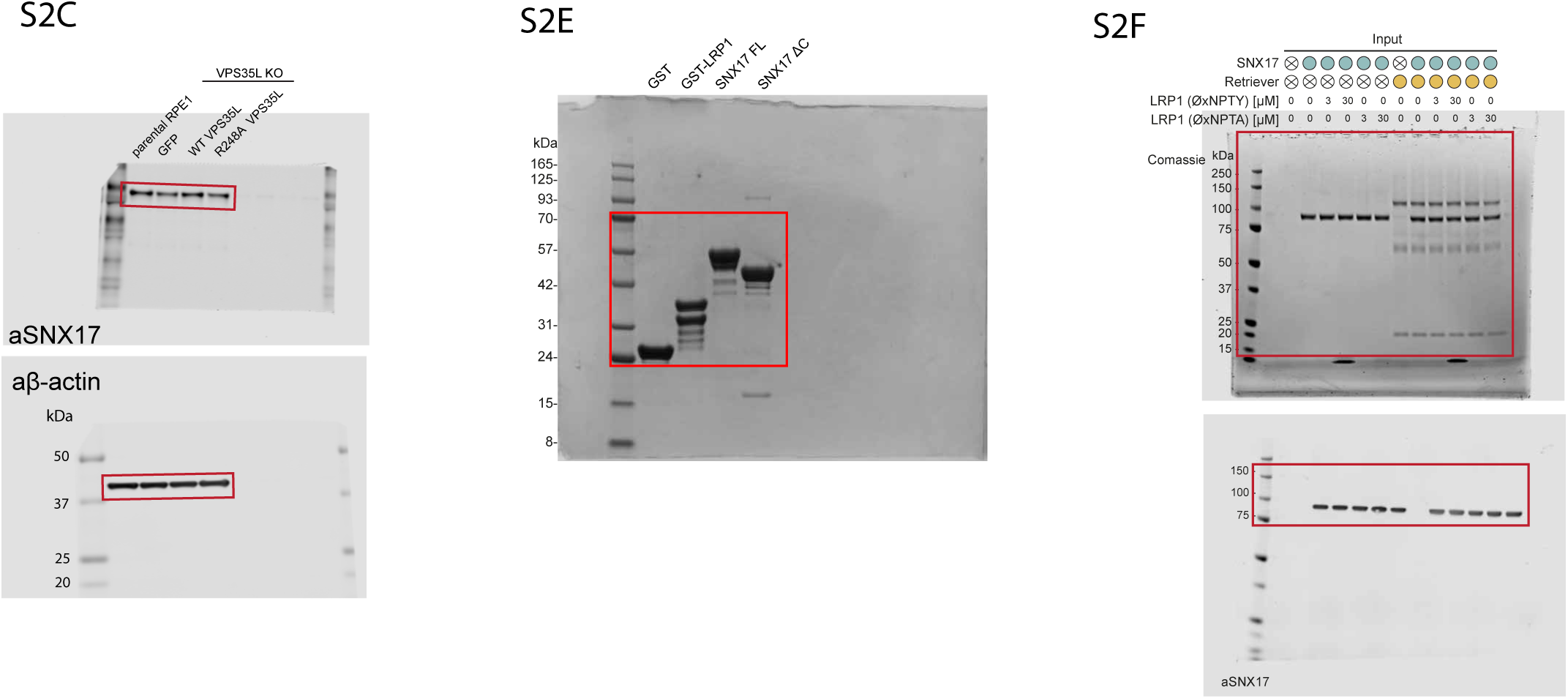

